# Improved identification of peptides, modification sites, and cross-link sites by Target-enhanced Accurate Inclusion Mass Screening (TAIMS)

**DOI:** 10.1101/2025.08.12.669802

**Authors:** Adalet Memetimin, Ching Tarn, Peng-Zhi Mao, Zhen-Lin Chen, Hao Chi, Yong Cao, Si-Min He, Meng-Qiu Dong

**Affiliations:** College of Life Sciences, Beijing Normal University, 19 Xinjiekouwai Avenue, Beijing 100875, China; National Institute of Biological Sciences, Beijing 102206, China; Tsinghua Institute of Multidisciplinary Biomedical Research, Tsinghua University, Beijing 100084, China; Key Laboratory of Intelligent Information Processing of Chinese Academy of Sciences (CAS), Institute of Computing Technology, CAS, Beijing, 100190, China; University of Chinese Academy of Sciences, Beijing, 100049, China

## Abstract

Chemical cross-linking of proteins coupled with mass spectrometry provides structural insights by identifying cross-linked peptide pairs, abbreviated as cross-links. Presently, cross-link identification suffers from ambiguity and poor sensitivity, because they are typically of lower abundance and consequently of lower MS2 quality than linear peptides present in the same sample. Here, we present Target-enhanced Accurate Inclusion Mass Screening (TAIMS), a meticulously optimized targeted mass spectrometry method. TAIMS significantly improved the quality of MS2 as indicated by fragment ion coverage (FIC) and other metrics. From data-dependent acquisition (DDA) to TAIMS, high-FIC cross-links increased by 401 or 678 on yeast ribosome or *E. coli* lysate, respectively, or from about 40% to around 90%. As a result, TAIMS substantially enhanced the accuracy of cross-link site localization and mitigated sensitivity loss in cross-link identification from large database searches. Enhanced identification sensitivity of TAIMS is further evidenced by its capacity to recover genuine cross-link identifications from data that would typically be discarded. On cross-linked *E. coli* lysate, 10.5% (284/2711) of the inclusion-list entries generated from unidentified cross-link-spectrum matches gained identity through TAIMS. Of these, 230 were linear peptides and 54 were cross-links, including 10 inter-molecular cross-links missed entirely by DDA. Additionally, we demonstrate that TAIMS is a general method for identification of low-abundance, post-translationally modified peptides. On a mouse brain sample, TAIMS increased the number of phosphopeptides identified with accurate phosphosite assignment by 67%. These findings indicate that TAIMS holds broad applicability in proteomics.

## Introduction

Chemical cross-linking of proteins coupled with mass spectrometry (abbreviated as CXMS, XL-MS or CLMS) is a convenient tool for probing protein structures and protein-protein interactions^1^. A successful CXMS experiment relies on multiple factors including cross-linker, prefractionation when it is needed, data acquisition, and data analysis.

Driven by the need to improve the sensitivity and efficiency of CXMS analysis, researchers have developed cross-linkers that are enrichable, gas phase cleavable, or both. Leiker^2^, Phox^3^, and tBu-PhoX^4^ are examples of enrichable cross-linkers; DSSO^5^, DSBU^6^, PIR^7^, BAMG^8^, and TDS^9^ are examples of cleavable cross-linkers; DSBSO^10^ and pBVS^11^ are both. Also explored were different chromatographic prefractionation methods, such as size exclusion chromatography^12,13^, strong cation exchange chromatography^14^, and hydrophilic strong anion exchange chromatography^15^. To bring out the best from gas phase cleavable cross-linkers, alternative data acquisition methods were adopted, including MS2-MS3 under collision-induced dissociation (CID)^16^, CID MS2 followed by electron-transfer dissociation MS2^17^, and MS2 using stepped higher-energy collisional dissociation (stepped HCD)^18,19^. For data analysis, a multitude of search engines have been developed for identification of cross-linked peptide pairs (abbreviated as cross-links). These include xQuest^14^, pLink^1,20^, Protein Prospector^21^, Xi^22^, Kojak^23^, Merox^24^, ECL^25^, XlinkX^16^, MetaMorpheusXL^26^, MS Annika^27^, MaxLynx^28^, CRIMP^29^ and Scout^30^. Additionally, several benchmark datasets were generated to evaluate the performance of search engines^15,30–32^. Through these continued efforts, CXMS has become an indispensable tool in structural biology.

Despite the progress made, the main challenge for CXMS persists, which is that cross-links are typically of low abundance^3^. This is exacerbated by intricate fragmentation patterns of cross-linked peptides and sometimes a huge search space arising from combining tens of thousands of peptide sequences or more. Under DDA — the ubiquitous data acquisition mode in CXMS experiments — low-abundance cross-links often suffer from insufficient ion accumulation. This leads to low-quality MS2 data, which in turn leads to low-confidence identifications or lack of identification. Improving the quality of MS2 is crucial for identifying more cross-links with higher confidence.

Targeted mass spectrometry is theoretically a good way to obtain high-quality MS2 for low-abundance ions. Targeted mass spectrometry has traditionally been employed mainly for quantitation during the validation phase, to complement DDA or data-independent acquisition (DIA)-based discovery workflows^33,34^. Recent intelligent acquisition strategies such as hybrid DIA^35^ and multiplexed DIA^36^ have demonstrated that integrating semi-targeted acquisition into DIA workflows can substantially enhance sensitivity and specificity in complex proteome analysis. Quantitation by targeted MS has evolved from selected reaction monitoring (SRM)^37,38^ to accurate inclusion mass screening (AIMS)^39–43^ and parallel reaction monitoring (PRM)^44–48^. The number of targets that can be covered in a typical targeted MS experiment has increased to over a thousand. The number of target ions and the amount of allowable time for each target, which directly affects MS2 quality, must be well balanced for optimal effect.

To this date, application of targeted MS to CXMS has been primarily for quantitation^49–50^, not identification. Here, to improve the quality of MS2 and subsequently to identify more cross-links with higher confidence, we tuned meticulously the parameters of AIMS and renamed the modified version Target-enhanced AIMS or TAIMS. In TAIMS, MS2 data acquisition is directed to candidate cross-links identified from DDA, with maximal ion injection time increased to 500 ms, AGC target raised to 2E5, resolution kept high at 120K, and dynamic exclusion time reduced to 3 seconds (using Fusion Lumos as an example). This markedly improved the quality of MS2 of low-abundance cross-links and increased high-confidence cross-link identifications at the precursor level by 53–131%. We demonstrate that TAIMS is a general method that can be applied readily to MS analysis of non-cross-linked peptides such as phosphopeptides, whose abundances vary over a vast dynamic range. Further, MS2 quality improvement brought about by TAIMS is highly beneficial for locating the exact sites of post-translational modification or cross-linking. To streamline the process of configuring a TAIMS data acquisition method as well as to analyze the data, we developed a software tool called TargetWizard and made it freely available.

## Results

### Optimization of MS parameters for targeted MS

Accurate and sensitive identification of cross-linked peptide pairs depends on the quality of MS2 spectra. We assessed how the MS instrument parameter settings in DDA, particularly those for MS2, affected the quality of MS2 of cross-links (Supplementary Figure 1). The quality of MS2 improved as the AGC target, the maximum ion injection time, and resolution increased, as indicated by better *E-*values of the CSMs (Supplementary Figure 1A-D). However, as a previous study shows, increasing ion injection time and resolution without considering acquisition speed is impractical in DDA, as it compromises the overall identification sensitivity^51^.

To balance spectral quality with acquisition speed, we shifted our attention from DDA to targeted mass spectrometry, which analyzes only predefined targets. Among the three primary targeted MS methods—SRM, AIMS, and PRM—we chose AIMS. Although AIMS and PRM both direct data acquisition towards precursors from an inclusion list, AIMS triggers MS2 only upon detecting a peak in the full MS scan that matches a listed *m/z* value, whereas triggering of MS2 in PRM is independent of the full MS scan. A key advantage of AIMS over PRM is its ability to analyze more peptides per MS run^40,52^.

AIMS has been employed in various applications, such as biomarker discovery^39,53,54^, quantitative proteomics^52,55^, and in-depth characterization of peptide mixtures^42^. In most of these studies, AIMS experiments were performed with DDA-like parameters (Figure 1A) and the improvement in MS2 quality over DDA is unremarkable. We thus investigated whether spectral quality could be improved categorically through rigorous optimization of the instrument parameters in AIMS experiments, and found that it is necessary to deviate from DDA-like settings.

**Figure 1.**
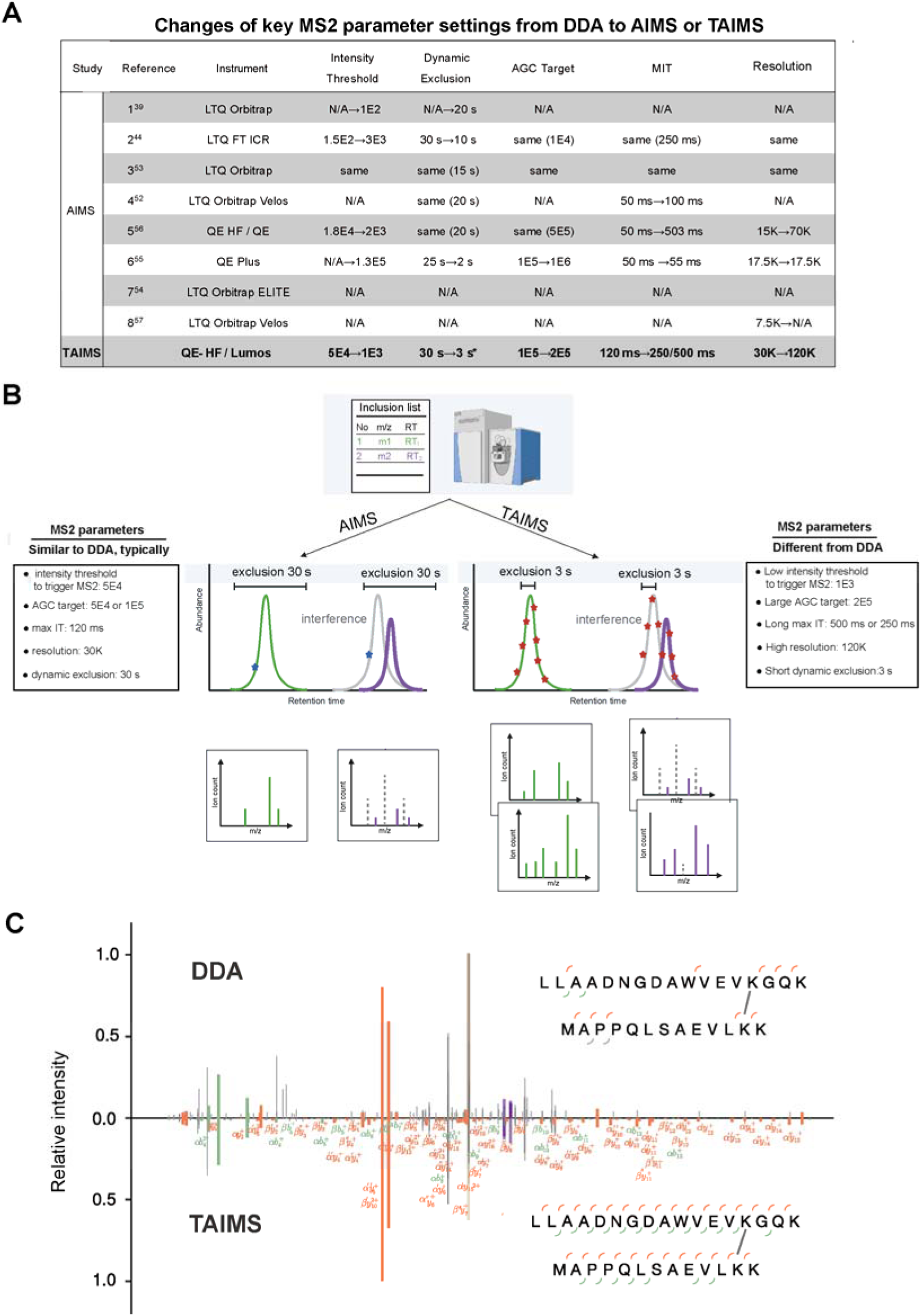
Development of TAIMS. **A** Key MS2 parameters from DDA to AIMS and TAIMS. “N/A” not available; "same" indicates that the parameter is identical for both DDA and AIMS methods; specific values are indicated if available. * A dynamic exclusion time of 3 s is optimal for this LC-MS run of a mean peak width of ∼20 s. In general, set dynamic exclusion time to 1/7–1/5 of the mean peak width (Supplementary Figure 1H), as determined from preceding DDA runs. **B** Like AIMS, TAIMS acquires MS2 spectra for targets of certain m/z, charge (z), retention time range as specified by an inclusion list. Compared with AIMS, TAIMS is characterized by shorter dynamic exclusion time, longer ion injection time, and higher resolution. As a result, TAIMS acquires more higher-quality MS2 spectra across the chromatographic peak of a target ion, which improves identifications of low-abundance ions in both quantity and quality. **C** DDA and TAIMS MS2 of a DSSO cross-link. Precursor ions are shown in light brown and DSSO doublets in purple. Long-arm and short-arm fragments are denoted by (′) and (″), respectively.

Using the optimal AGC and resolution settings obtained from DDA (Supplementary Figure 1C-D), we further optimized the maximum ion injection time (MIT) and the dynamic exclusion time for AIMS using a disuccinimidyl sulfoxide (DSSO) cross-linked bovine serum albumin (BSA) sample (Supplementary Figure 1E-G). This set of experiments revealed one striking difference in dynamic exclusion time between DDA and AIMS: whereas 15–30 s is the norm for DDA (Figure 1A), setting it to zero seconds resulted in more cross-link identifications than 10 s and 30 s under AIMS (Supplementary Figure 1E). After further refinement with parallelizable injection time taken into consideration, we found that a dynamic exclusion time of 3 s is optimal for AIMS in our liquid chromatography-mass spectrometry (LC-MS) runs in which the mean peak width was 20 seconds (Supplementary Figure 1E-G). We varied the length of LC-MS runs and found that setting dynamic exclusion to 1/10 - 1/5 of the mean peak width is recommendable for AIMS experiments (Supplementary Figure 1H).

Lowering the global intensity threshold level from 5E4 to 1E4 or 1E3 boosted MS2 acquisition for low-intensity precursor ions (Supplementary Figure 1I). In the inclusion-list window of Orbitrap Fusion Lumos, intensity threshold can be specified for individual precursor ions—*i.e.*, individualized intensity threshold (IIT)—on top of a global intensity threshold. We observed that enabling IIT at a level of 10-25% increased slightly the number of cross-link identifications from yeast ribosomes treated with DSSO (Supplementary Figure 1J).

The optimized parameter settings for AIMS are summarized in Figure 1A (last row). For MIT, two options are suggested: 250 ms and 500 ms, approximating two parallelizable injection time values of 246 ms and 502 ms (Supplementary Figure 1F), respectively. MIT of 500 ms is recommended for an inclusion list of 2000 target ions or less, whereas 250 ms is recommended for an inclusion list of 2000∼4000 target ions. Not included in Figure 1A is an optional IIT setting at 10%; setting it up adds an extra layer of complexity and the improvement is marginal (Supplementary Figure 1J). Lastly, using the optimized parameters, a side-by-side comparison of AIMS and PRM was conducted. As shown, AIMS consistently outperformed PRM (Supplementary Figure 1K).

This optimized AIMS method fully utilizes the MS2 data acquisition time on predefined targets. It ensures sufficient accumulation and repetitive acquisitions for each target ion in the predefined inclusion list (Figure 1B). We named this modified AIMS as target-enhanced AIMS, or TAIMS (Figure 1). Representative spectra from DDA and TAIMS are shown in Figure 1C.

### TAIMS markedly improves MS2 quality

A workflow for TAIMS analysis following initial DDA was established for the identification of cross-linked peptides, as illustrated in Figure 2. To quantify the spectral improvement provided by TAIMS over DDA, a DSSO cross-linked *E. coli* whole-cell lysate was prepared. DDA analysis of this sample, searched with pLink 2.4.2 using a 5% precursor FDR, identified 1299 cross-linked peptide precursors. To further obtain candidate cross-links, a preliminary search with a relaxed 100% precursor FDR was performed, resulting in 4010 cross-linked peptide precursors.

**Figure 2.**
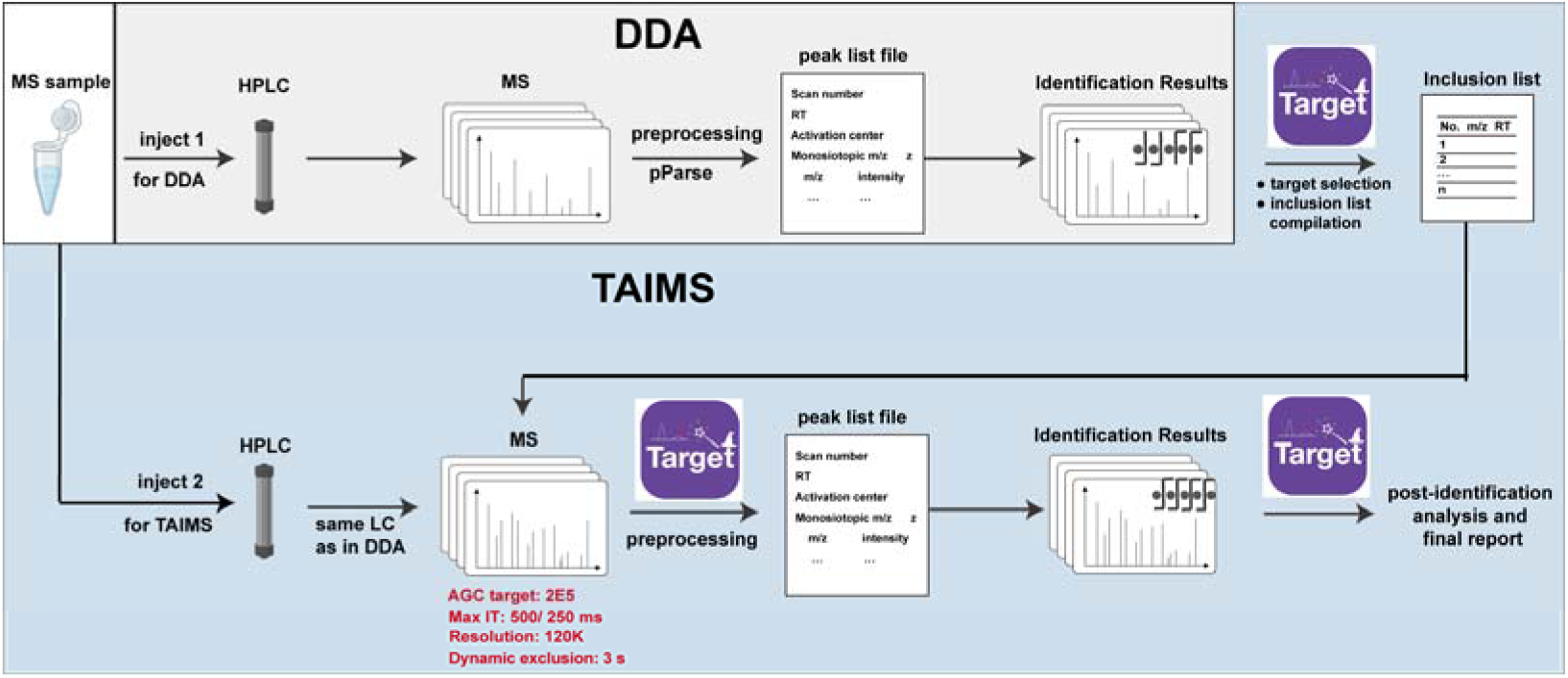
A generalized TAIMS workflow for cross-linked peptides. Cross-linked samples are first analyzed using DDA and searched with pLink 2.4.2. Precursor ions corresponding to identified and candidate cross-links from the DDA run are re-analyzed using TAIMS to identify more cross-links at higher confidence levels.

These comprised 1299 identified cross-links (as noted above) and 2711 candidate cross-links, with 841 matched to the target database but not passing the 5% FDR cutoff, and 1870 matched to the decoy database (Figure 3A). Precursor ions from all three groups were extracted for TAIMS analysis.

**Figure 3.**
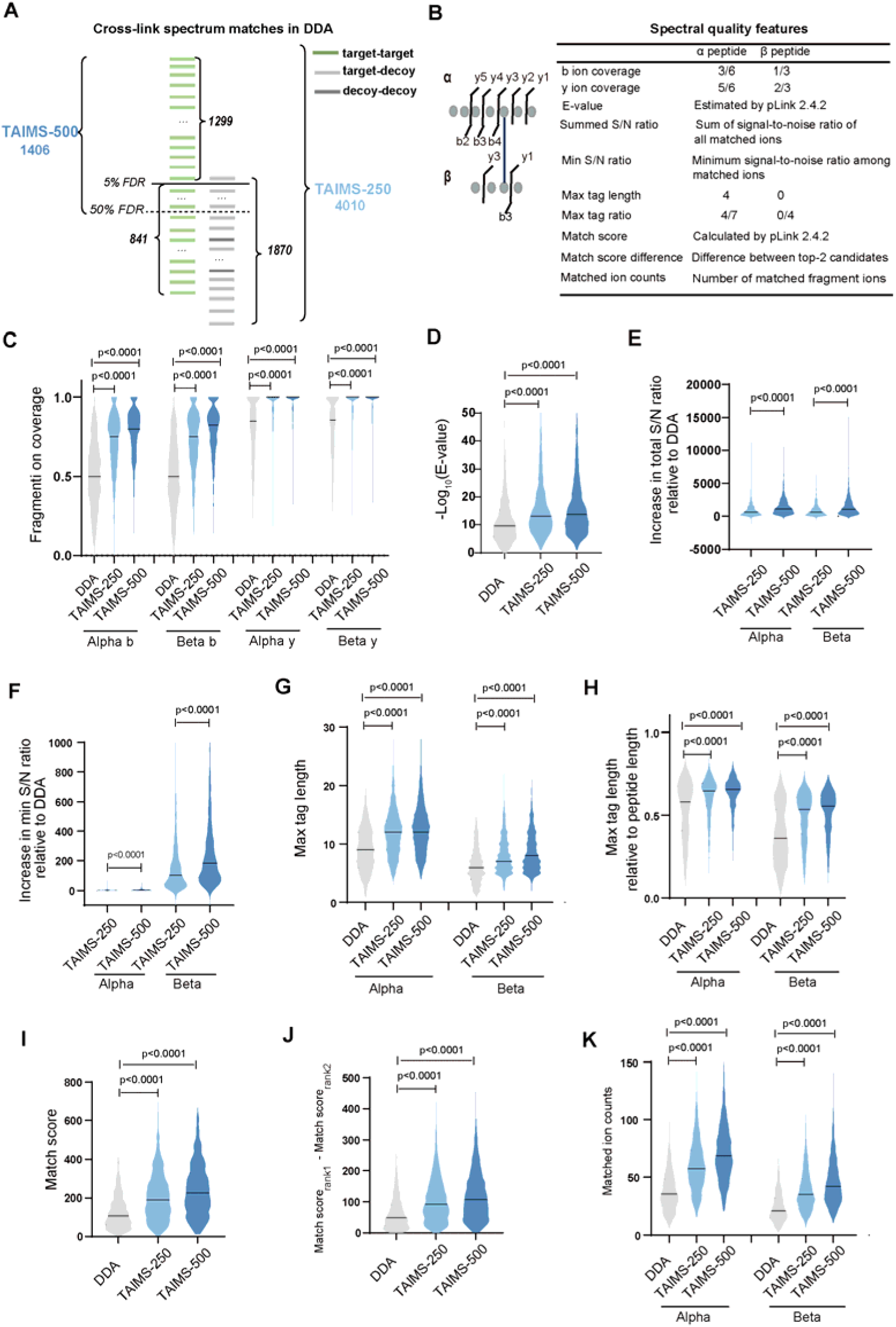
TAIMS MS2 spectra are of higher quality than DDA MS2 spectra. **A** Selection of target precursor ions for TAIMS. The DDA data of a DSSO cross-linked *E. coli* lysate was searched using pLink2 and filtered with a 5% FDR at the level of precursor ions. Two inclusion lists were compiled as indicated, one for TAIMS-250 (MIT 246 ms) and the other for TAIMS-500 (MIT 502 ms). **B** Nine spectral quality metrics. **C-K** Comparison of 1175 sets of cognate DDA, TAIMS-250, and TAIMS-500 MS2 spectra. A common set of 1175 precursor ions of cross-linked peptide pairs were identified from a DSSO cross-linked *E. coli* lysate sample across three MS runs, one DDA and two TAIMS. To compare among the cognate MS2 spectra, CSMs with the best *E*-values were chosen, one each from DDA, TAIMS-250, and TAIMS-500. For pLink2 search of the DDA and the TAIMS data, fragment ion mass tolerance was set to ±20 ppm and ±5 ppm, respectively. For statistical analysis of the identified DDA and TAIMS spectra, a single fragment ion mass tolerance of ±20 ppm was used. Alteration of this to ± 20 ppm for DDA MS2 and ±5 ppm for TAIMS MS2 made little difference (Supplementary Figure 2). Of these, max tag length (**G**) is the maximum length of consecutive *b* or *y* fragment ions; for a precursor ion, the match score (**I**) is the fine score of the rank-1 candidate of the best matched spectrum according to the pLink2 search result; match score difference (**J**) is the fine score difference between rank-1 and rank-2 candidates. Statistical significance was assessed using the Wilcoxon matched-pairs test.

Two TAIMS runs—TAIMS-250 and TAIMS-500—were conducted following the initial DDA run (Figure 3A). The inclusion list of TAIMS-250 included all 4010 precursor ions (1299+2711) for a comprehensive reanalysis. In contrast, TAIMS-500 focused on 1406 precursor ions that passed a 50% precursor FDR cutoff and ignored those that did not. The inclusion lists were generated using TargetWizard, a software tool that we developed to streamline method setup and data reporting for TAIMS experiments. Free download of TargetWizard is available at https://targetwizard.ctarn.io.

To compare TAIMS MS2 and DDA MS2 of the same target ion, we began data analysis by registering TAIMS MS2 spectra with their intended target ions. An MS2 spectrum that conforms to the specifications of precursor m/z, charge (z), and retention time is not necessarily an MS2 spectrum of the intended target ion, so we implemented a process to discriminate cognate MS2 spectra from unrelated ones. Details are provided in Methods. Briefly, each TAIMS MS2 is logged under a target ion if their *m/z*, z, and retention time values all aligned (*m/z* tolerance ±10 ppm, exact match for charge state and retention time window). Subsequently, the DDA MS2 of a target ion is compared to each TAIMS MS2 logged under the same target ion. If the spectral similarity score exceeds a threshold determined by a discrimination model trained on-site, the TAIMS MS2 is classified as cognate, which indicates successful triggering of an MS2 scan for the intended target ion. In this way, we determine that in TAIMS-250 and TAIMS-500 runs, MS2 scans were triggered successfully for 97% (3900 out of 4010) and 99% (1396 out of 1406) of the precursor ions in the inclusion list, respectively.

Using the above data, we evaluated the quality of MS2 spectra of TAIMS-250 and TAIMS-500 runs against their cognate DDA MS2 spectra (Figure 3). A total of nine metrics were used to assess quality of MS2 spectra, including *b/y* fragment ion coverage, *E*-value, summed signal-to-noise ratio, minimum signal-to-noise ratio, maximum tag length, maximum tag ratio, match score, match score difference, matched fragment count (Figure 3B). Since these are MS2 of cross-linked peptide pairs, we evaluated the α peptide (of a relatively higher mass) and the β peptide (of a relatively lower mass) separately. As shown in Figure 3C-K, TAIMS MS2 spectra are markedly and consistently better than DDA MS2 spectra across all nine spectral quality metrics. The TAIMS-500 spectra are slightly better than or on par with the TAIMS-250 spectra. The improvement in MS2 quality with TAIMS is primarily attributed to the “enrichment” of target ions in MS2 scans (Supplementary Figure 3). This advantage of TAIMS is further validated using DSS (disuccinimidyl suberate) as the cross-linking reagent (Supplementary Figure 4).

### TAIMS boosts the precision of cross-link identification

We investigated how the enhancement of MS2 quality by TAIMS translates into improved cross-link identification. Given that not all cross-link identifications are equally reliable, there is a need to distinguish reliable ones from those that are not. As shown in Supplementary Figure 5, *b/y* fragment ion coverage provides a useful distinction between correct and incorrect identifications. This provides a basis on which a set of stringent criteria is established for high-quality cross-link identification: For both the alpha peptide and the beta peptide, the *y*-ion coverages must exceed 0.6, and the *b*-ion coverages must exceed 0.4.

Next, we assessed the quality of identification of 1299 cross-link precursor ions from the DDA analysis of DSSO-linked *E. coli* lysate and found that 43.3% (563/1299) of them were high-FIC identifications. After re-analysis by TAIMS, 87.6% (1138/1299, by TAIMS-250) and 95.5% (1241/1299, by TAIMS-500) of them became high-FIC identifications (Figure 4A). Figure 4B shows that the sequence coverage of the alpha peptide and the beta peptide by fragment ions is significantly higher in the TAIMS MS2 data compared to the DDA MS2 data, further confirming the accuracy of the original identifications. Furthermore, TAIMS pinpoints a false DDA identification (Supplementary Figure 6).

**Figure 4.**
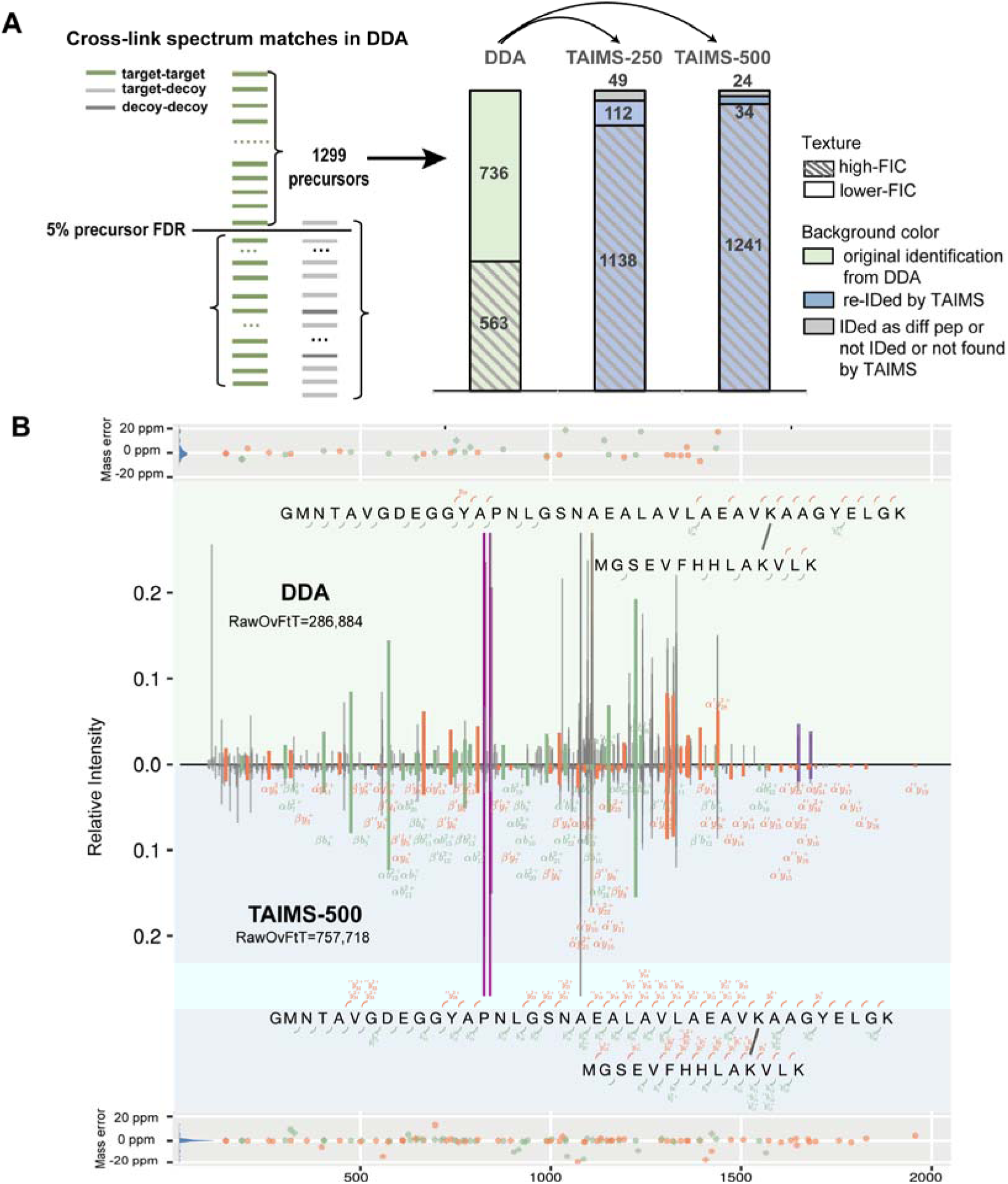
TAIMS improves cross-link identification precision. **A.** TAIMS analysis outcome of cross-link precursor ions identified in DDA. **B** Example of MS2 quality improvement in TAIMS. Of all the fragment ions, only the ones that were detected in TAIMS but not DDA, or in DDA but not TAIMS are labeled. Total ion count of a MS2 scan was recorded in the raw file as a parameter “RawOvFtT”. Note that the total ion count of the TAIMS MS2 more than doubled compared to that of the DDA MS2 of the same precursor.

DDA-TAIMS analysis of a DSSO cross-linked yeast ribosome sample produced similar results (Supplementary Figure 7).

These results demonstrate that TAIMS re-analysis of precursor ions identified by DDA can provide the following benefits: (1) confirmation of identification, (2) elevation of low-FIC identification to high-FIC identification, and (3) revelation of false or less reliable identifications.

### TAIMS Identifies additional cross-links from DDA precursors of subthreshold CSMs

DDA analysis of the cross-linked *E. coli* lysate generated 2711 candidate cross-links, comprising 841 CSMs below 5% FDR and 1870 in the decoy space (Figure 5A). In an attempt to find treasure from the trash, an inclusion list was built from 2711 precursors corresponding to these unidentified CSMs. In the resulting TAIMS-250 run, 284 (10.5%) of these precursors were identified. Most (230/284) corresponded to mono-links, loop-links or linear peptides not modified by DSSO, but 54 were cross-links (Figure 5A). Among these, 41 were not identified in the initial DDA run.

**Figure 5.**
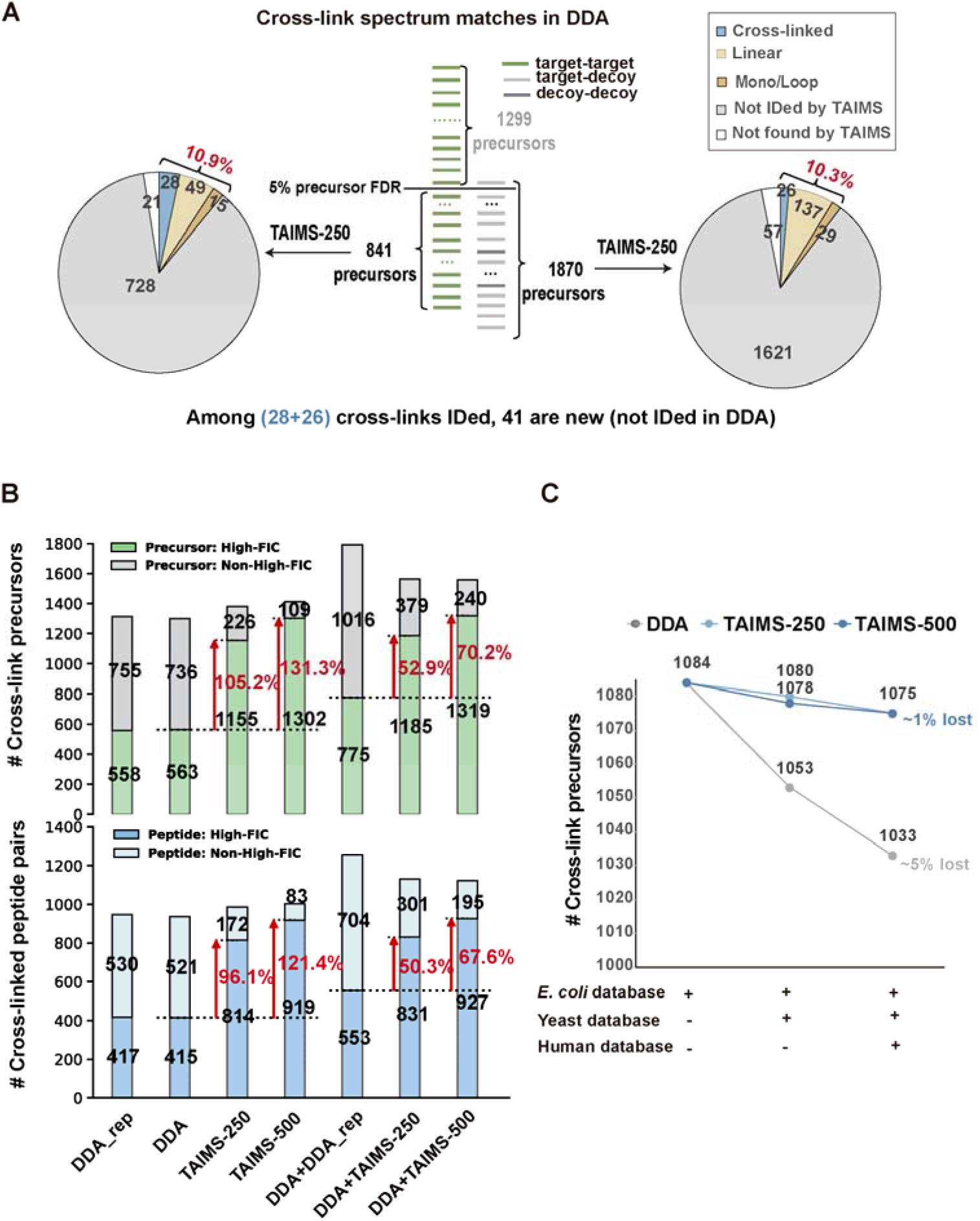
TAIMS improves cross-link identification sensitivity**. A** TAIMS analysis gained identity for 10.5% of the targets with unidentified DDA CSMs. **B** Overall increase of high-FIC cross-link identifications in TAIMS. The graph compares the number of identified cross-link precursors or cross-linked peptide pairs, both with and without the high-FIC requirement. A mass tolerance of ±20 ppm was applied for FIC calculation. **C** TAIMS spectra are resistant to sensitivity loss in cross-link identification caused by search space expansion. The DDA, TAIMS-250, and TAIMS-500 data of the DSSO cross-linked *E. coli* lysate sample yielded a common set of 1084 precursors when searched against the *E. coli* database. The corresponding DDA spectra, but not the TAIMS spectra, suffered considerable sensitivity loss when searched against expanded databases that included the yeast and the human databases in addition to the *E. coli* database.

To investigate why TAIMS gained identity for only 284 (10.5%) of the 2711 precursors on the inclusion list, we analyzed the corresponding MS1 spectra in the DDA data. This examination revealed that 1354 of them were dubious targets without experimental evidence supporting their assigned monoisotopic m/z and charge (Supplementary Figure 8A). This issue arises because the MS-raw-data-preprocessing software outputs one or more sets of monoisotopic m/z and z for each MS2 scan, and these candidate precursor or precursors will have a chance to explain the fragment ions detected in the MS2 scan. Preprocessing is not error-proof; in fact, it tends to err on the side of retaining all potential candidates. Therefore, some of the extracted MS2 scans from DDA are paired with false or “imagined” monoisotopic m/z and z values.

These imagined precursors may be selected for targeted MS analysis (Supplementary Figure 8B). To search TAIMS data, TargetWizard pairs each TAIMS MS2 with the trigger m/z and z specified in the inclusion list. Naturally, TAIMS MS2 of imagined precursors are destined to fail in identification due to incorrect precursor mass assignments.

Of the 41 newly identified cross-links by TAIMS, 10 were inter-protein ones. This is not insignificant, given that only 25 inter-protein cross-links were identified initially by DDA. Of the ten “salvaged” inter-protein cross-links, eight originated from the group of 841 precursors (target matches, FDR > 5%), one from the group of 1870 (decoy matches), and one from the group of 1299 (target matches, FDR ≤ 5%). Supplementary Figure 9 illustrates an inter-protein cross-link missed by DDA (q-value > 0.7) but confidently identified by TAIMS-250.

Similar results were observed in the TAIMS analysis of unidentified DDA CSMs from a cross-linked yeast ribosome sample (Supplementary Figure 7).

Taken together, by targeting precursor ions initially identified or not, TAIMS achieved a modest increase in the total number of cross-link identifications but led to a substantial rise in high-FIC identifications (Figure 5B). Compared to a single DDA run, DDA followed by TAIMS-250 or TAIMS-500 increased the number of high-FIC cross-link precursor identifications by 105.2% or 131.3%, respectively. Compared to two DDA runs, the increase was 52.9% or 70.2%, respectively (Figure 5B, upper panel). Similar trends were observed at the cross-linked peptide pair level (Figure 5B, lower panel).

In the late stage of this study, pLink 3 became available. To address a concern that the benefit of TAIMS might dissipate upon the advent of a more powerful search engine, we repeated the entire DDA–TAIMS workflow on the cross-linked yeast ribosome sample using pLink 3.0.17. This experiment demonstrated that the improvement by TAIMS is comparable using either pLink 2.4.2 or pLink 3.0.17 (Supplementary Figure 10).

### TAIMS alleviates sensitivity loss in proteome-wide cross-link search

Cross-link identifications from complex samples such as cell lysates suffer a great deal of sensitivity loss, particularly for inter-protein cross-links. This is, at least in part, because the search space for inter-protein cross-links scales quadratically with the number of protein sequences in the database^58,59^. MS2 quality is another factor that affects the sensitivity of cross-link identification. Evaluations conducted on simulated, high-quality cross-link spectra have shown that pLink1 and pLink2 can maintain a sensitivity level above 99% when searched against a human entrapment database^20^. We thus reasoned that high-quality MS2 spectra obtained through TAIMS should effectively mitigate sensitivity loss in a proteome-wide cross-link search.

As shown, of the *E. coli* cross-links identified from both the DDA and the TAIMS data, the quality of MS2 is much higher in the TAIMS data than in the DDA data (Figure 3). As expected, the high-quality TAIMS data are resistant to sensitivity loss when challenged by search space explosion, losing no more than 10 cross-links from a total of 1084 (Figure 5C). In comparison, DDA data suffered a loss of up to 51 cross-links under the same conditions (Figure 5C). MS2 spectra are shown for one such cross-link (Supplementary Figure 11). The same pattern was observed in a DSSO cross-linked yeast ribosome sample (Supplementary Figure 12).

We further analyzed the DDA CSMs that were lost in large database search. Half of the MS2 spectra were assigned to sequences in the reversed databases or the entrapment databases. The other half, although matched to the original, correct sequence pairs, suffered elevation in *q*-value great enough to be filtered out. Both scenarios are caused by increased interference associated with search space expansion. Taken together, TAIMS spectra, due to their higher fragment ion coverage over DDA spectra, effectively mitigate sensitivity loss in cross-link search at the systems level, *e.g.,* in protein-protein interactome analysis.

### Prioritizing DDA targets for TAIMS analysis

A TAIMS experiment has a finite capacity. The exact number of target precursor ions at full capacity is determined by the LC-MS/MS condition. In our experiments (see Methods), the inclusion list can have about 2000 precursor ions for TAIMS-500 or 4000 for TAIMS-250. If a DDA-derived target list is too long to fit into a single TAIMS run, either the list is shortened by prioritizing the targets or more than one TAIMS run arranged, or both.

Before we look for ways to shorten an inclusion list, let us be clear about the purpose of TAIMS. We envision that for cross-links that have passed a certain FDR threshold, i.e., the identified ones, TAIMS serves to increase high FIC identifications and to remove false positives as much as possible. Secondly, TAIMS serves to remove false negatives by salvaging true cross-links from the unidentified ones (e.g., FDR >5%).

It is obvious that high-FIC DDA identifications can be the first ones to go off the inclusion list, and this amounts to about 40% of the identified (Figure 4A and Supplementary Figure 7). To find out other ways to trim the list, we performed ROC analysis on multiple candidate metrics (Supplementary Figure 13A) and found that SVM score and *E-value* outperformed others. For DDA identifications sorted by either SVM score or *E-value*, the ones unconfirmed by TAIMS — with higher probability of being false—are concentrated in the worse 50%, although a single unconfirmed precursor is seen at the 30th percentile (Supplementary Figure 13B–C).

This means that the better 30% or 50% need not be re-analyzed by TAIMS because it is unlikely to find false positives among them. Alternatively, DDA identifications with *E-value* <1×10^−8^ or more stringently, *E-*value <1×10^−1^^3^ need not be included (Supplementary Figure 13C). Similarly, for DDA CSMs that fail to pass the FDR cutoff—the unidentified ones—those that are salvaged by TAIMS are concentrated in the top 20% by SVM score or FDR (Supplementary Figure 13D-F, sorted after CSMs of poor isotopic evidence are removed). *E-value* was not included here because pLink does not output it for unidentified CSMs. In brief, the bottom 80% of unidentified DDA CSMs are not worthy of re-analysis by TAIMS. Similar conclusions can be drawn from the yeast ribosome data (Supplementary Table 1).

If a trimmed inclusion list still exceeds the capacity of a single TAIMS run, TargetWizard supports multi-batch scheduling, in which precursor ions are divided into two or more LC-MS/MS runs (Supplementary Figure 14).

### Precise localization of cross-link sites through TAIMS

Recently, the precision of cross-linked sites has been called into question^57^. Since TAIMS MS2 spectra tend to have high fragment ion coverage, we investigated whether this would facilitate locating the site of cross-linking. We define a cross-link site-associated fragment ion (site-FI) as an *a-*, or *b-*, or *y-*ion that is generated from a cleavage event occurring no more than 2-aa away from the cross-linked residue. Site-FI coverage, that is, site-FIC, is calculated as the count ratio between the observed site-FIs (20 ppm mass tolerance) and all theoretical site-FIs (Figure 6A).

**Figure 6.**
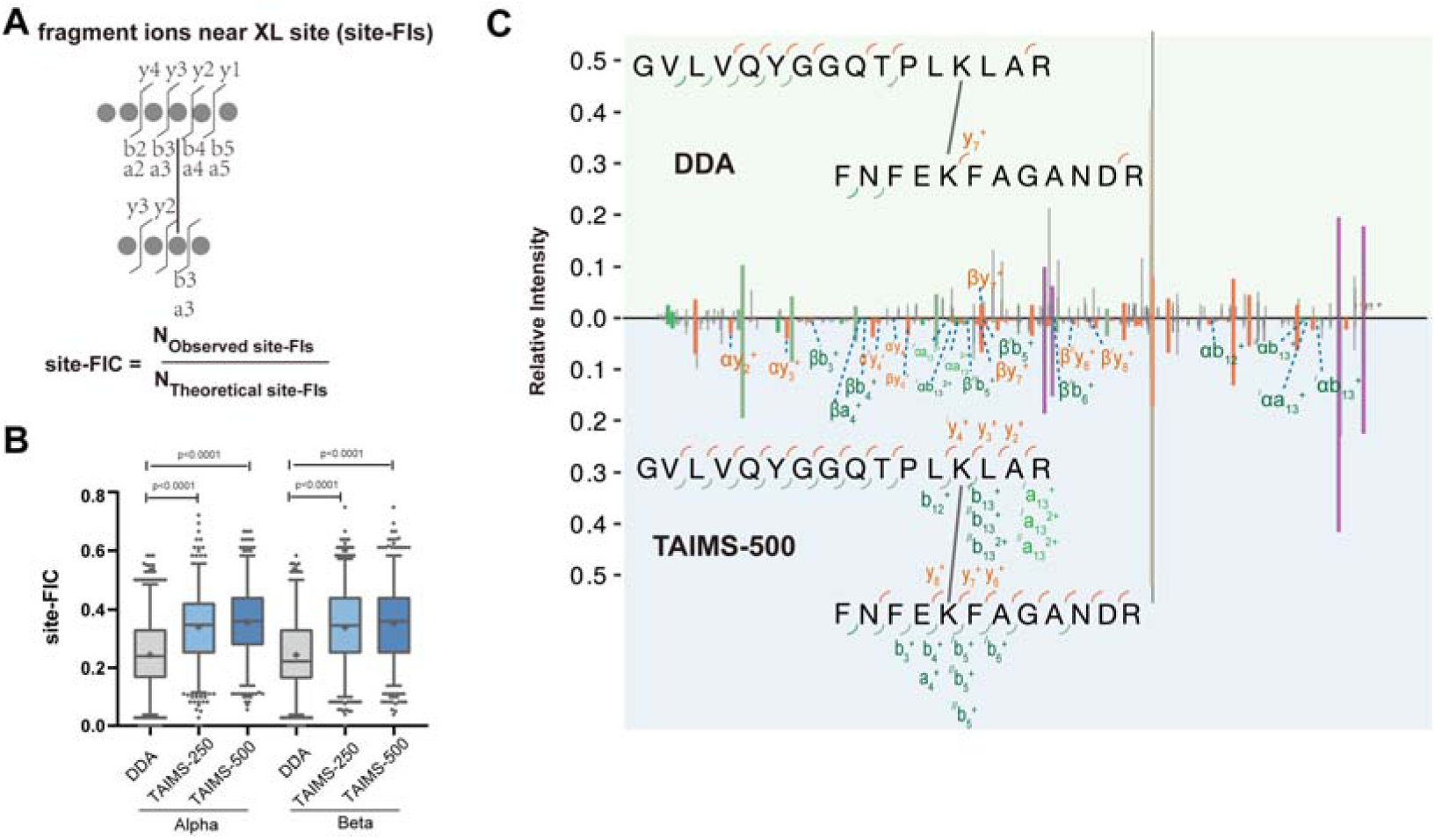
TAIMS improves the precision of locating cross-linked sites. **A** Site-FIC is the count ratio of observed over all theoretical a/b/y fragment ions generated no more than 2-aa away from the cross-linked residue. Only fragment ions with charge states of 1+ and 2+ are considered. In the example provided, all the theoretical fragment ions used for calculating site-FIC are enumerated. **B** Calculated site-FIC values for K-K cross-link precursor ions identified from DDA or TAIMS data based on the shared identifications from the cross-linked *E. coli* lysate sample. Maximum allowed mass deviation is 20 ppm between an observed and a theoretical fragment ion. Statistical significance was assessed using Wilcoxon matched-pairs test. **C** DDA and TAIMS MS2 identified a cross-link between the same two peptides with the same cross-linked sites. For clarity, labeling of fragment ions is limited to the site-FIs of two peptides.

Of the overlapped K-K cross-links identified from *E. coli* lysate by DDA, TAIMS-250 and TAIMS-500, the site-FIC values grouped by MS method, clustered around 0.24/0.22, 0.35/0.35, and 0.36/0.36, respectively, for alpha/beta peptide (Figure 6B). Hence, cross-link sites are mapped more precisely by TAIMS than DDA. As illustrated in Figure 6C, while the cross-link site localization remained consistent between DDA and TAIMS-500, the site-FIC values increased from 0.00/0.03 for alpha/beta-peptide in DDA to 0.31/0.25 in TAIMS-500.

Enhanced precision makes determination of cross-linked sites more accurate. One example is shown in Supplementary Figure 15. For the indicated cross-linked peptide pairs, the site-FIC values of the two possible cross-linked lysine residues (K52 and K60 of protein L7/L12) in the alpha peptide are 0.06 and 0.08, respectively, according to the DDA spectrum, or 0.06 and 0.19, respectively, according to the TAIMS spectrum. The cross-linked site in the alpha peptide was assigned to K60 based on strong evidence (αy7–αy13) in the TAIMS spectrum (Supplementary Figure 15, lower panel). It was assigned to K52 based on sparse and spurious evidence in the DDA spectrum (Supplementary Figure 15, upper panel). Note that two out of the three fragment ions (αb25++, αy14+, αy14++) that could support the K52 assignment have relatively large mass errors close to 10 ppm (Supplementary Figure 15, marked by colored dots in the mass error bar above the DDA MS2 spectrum).

NHS ester cross-links through serine (S), threonine (T), or tyrosine (Y) residues have recently been called into question, as statistical analysis results suggest that most STY cross-links identified should be false^60^. However, the percentage of false identifications was not estimated^60^. The MS2 data analyzed in this study are all acquired in the DDA mode. Since TAIMS spectra provide higher FIC and site-FIC, we wondered whether TAIMS would allow estimation of the exact fraction of probable versus improbable STY cross-links.

With STY considered as cross-linkable sites in addition to K and the protein N-terminus, pLink2 search retrieved a total of 171 precursor ions of STY cross-links from the TAIMS-500 data of the *E. coli* sample. As shown in Figure 7A, 15 of them corresponded to high-FIC cross-links (FIC > 0.6 by *y-*ions and FIC > 0.4 by *b-*ions for both the alpha peptide and the beta peptide) lacking a cross-linkable K. An additional 14 precursor ions corresponded to high-FIC cross-links where the purportedly cross-linked STY residue, although accompanied by a cross-linkable K in the same peptide, has more site-discriminating fragment ions (defined in Figure 7B) than the alternative, cross-linkable K. We regard these two groups of STY cross-links as probable and thus estimate that approximately 17% (15/171 + 14/171) of the STY cross-links identified are likely correct.

**Figure 7.**
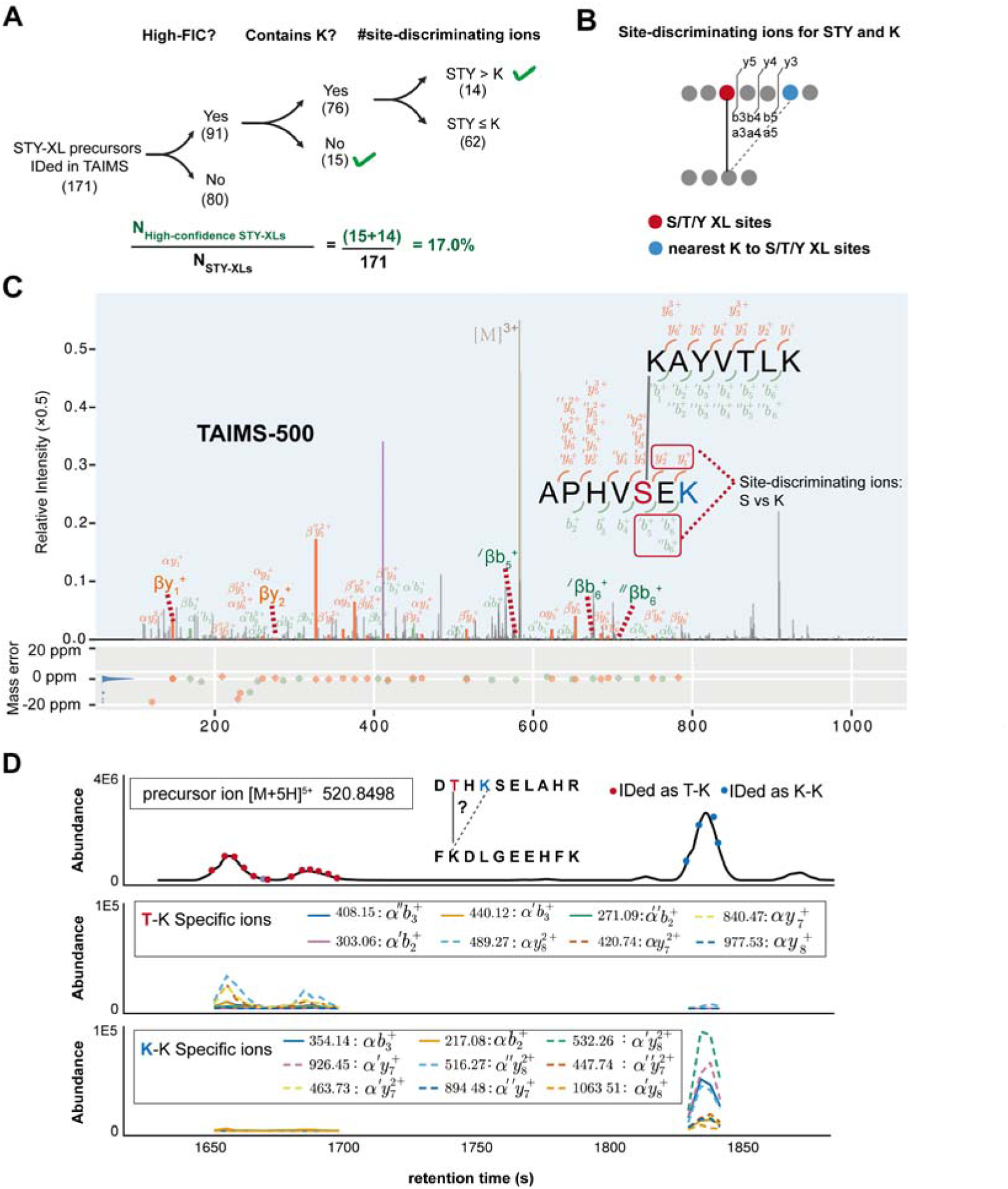
Estimation of the proportion of probable STY cross-links using data acquired by TAIMS. **A** Breakdown of the STY cross-links identified from a DSSO cross-linked *E. coli* sample. **B** Definition of site-discriminating fragment ions for two alternative cross-link sites, STY vs. K. **C** MS2 of an identified STY cross-link. **D** A T-K cross-link and its isoform of K-K cross-link elute at different times. Displayed are the reconstructed chromatograms of the precursor ions (top) and the fragment ions unique to the T-K (middle) or K-K (bottom) cross-link.

For one of these probable STY cross-links <u>K</u>AYVTLK-APHV<u>S</u>EK, its MS2 spectrum is displayed in Figure 7C. The site of cross-linking in the β peptide is assigned to a serine residue. The lysine residue at the C-terminus of the β peptide must not be the site of cross-linking because if it were, it could not have been cleaved by trypsin to produce this peptide^60^. Nevertheless, if assumed that this lysine residue could be cross-linked, there are no site-discriminating fragment ions for this lysine. In contrast, there are five site-discriminating fragment ions for serine (βy1□, βy2□, βLb5□, βLb6□, and βSb6□), as indicated by red dotted lines.

We also found an STY cross-link D<u>T</u>HKSELAHR-F<u>K</u>DLGEEHFK with an isoform of K-K cross-link DTH<u>K</u>SELAHR-F<u>K</u>DLGEEHFK (Figure 7D). Reconstructed chromatograms of fragment ions unique to the T-K cross-link or the K-K cross-link revealed that the T-K isoform has a shorter retention time than the K-K isoform. With the knowledge of the respective retention times and the reconstructed chromatogram of the precursor ions, we estimated that the T-K isoform and the K-K isoform have an abundance ratio of roughly 1:3.

Taken together, improved MS2 quality via TAIMS sharpened our understanding of STY cross-links: They do exist, but in small numbers. This is in line with conclusion of previous study^60^, but here we present the first quantitative estimate that approximately 17% of STY cross-links identified by pLink2 in large-scale datasets are likely correct.

### Precise localization of phosphorylation sites through TAIMS

Post-translational modification (PTM) analysis is integral to MS-based proteomics. Precise localization of a PTM site is necessary for understanding the biology of the PTM and requires high peptide sequence coverage by fragment ions. Here, we investigated whether improvement in MS2 quality through TAIMS facilitates PTM localization.

Phosphorylation is a common PTM essential for numerous cellular functions. To evaluate the performance of TAIMS in phosphorylation site localization, we analyzed a whole-mouse-brain phosphoproteomics sample. For 1567 out of the 5198 precursor ions of phosphorylated peptides identified by pFind3^61,62^ (FDR <1%), their phosphorylation site assignment is more likely wrong than correct (localization probability < 50% as determined by PTMiner^63,64^). We subjected this group of precursor ions to TAIMS analysis, along with 637 unidentified but perhaps “salvageable” precursor ions whose phosphopeptide-spectrum matches were in the FDR range of 1–10% (Figure 8A).

**Figure 8.**
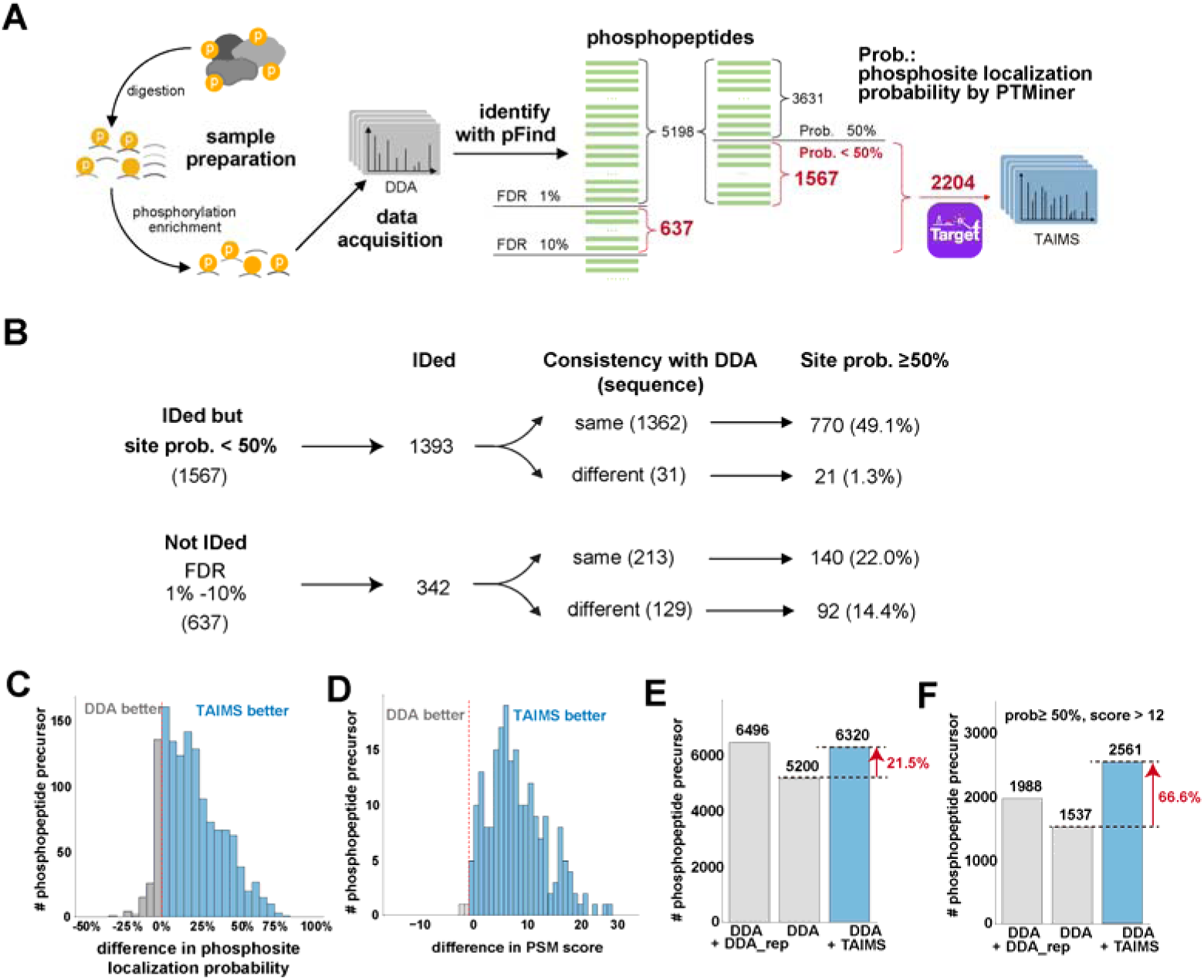
TAIMS enhances phosphorylated peptide localization precision. **A** Workflow for DDA-TAIMS analysis of phosphopeptides. Phosphopeptides were identified using pFind3 and phosphosite localization probabilities were calculated using PTMiner. **B** Result of TAIMS analysis. **C** Difference in phosphosite localization probability for the overlap of 1362 precursor ions of phosphopeptides identified in DDA and TAIMS data. **D** Difference in the pFind3 raw scores of 213 precursor ions whose DDA and TAIMS spectra matched the same phosphopeptides, but the DDA PSMs did not pass the 1% FDR cutoff and the TAIMS PSMs did. **E** Comparison of the identification results of DDA, DDA-DDA, and DDA-TAIMS. DDA_rep is a replicate DDA run, and DDA+DDA_rep represents a combined analysis of two independent DDA injections. No confidence filter was applied. **F** Similar to E, but with two confidence filters applied: phosphosite localization probability ≥ 50% and pFind3 PSM score > 12.

For the 1567 targets in the first group, 1362 had TAIMS spectra matched to the same phosphopeptides as their DDA spectra, hence validating the sequence identification. Of these, 770 had TAIMS spectra that provided a phosphosite localization probability ≥ 50% (Figure 8B). Another 21 targets gained a different identity through TAIMS, but each with a phosphosite localization probability ≥ 50% (Figure 8B). In total, TAIMS lifted 50.5% (770/1567 + 21/1567) of these targets out of the group of questionable phosphosites (i.e., localization probability increased from < 50% to ≥ 50%). As shown in Figure 8C, for the 85.7% (1167/1362) of the phosphopeptides identified by both TAIMS and DDA, TAIMS provided higher phosphosite localization probability.

After TAIMS analysis of the second group, which consisted of 637 targets whose DDA peptide-spectrum matches (PSMs) were unidentified, 232 (140 + 92) or 36.4% of them were identified as phosphopeptides with phosphosite localization probability greater than 50% (Figure 8B). Figure 8D displays the difference in the pFind3 PSM scores of 213 precursor ions that were assigned to the same phosphopeptides but only the TAIMS PSMs passed the 1% FDR cutoff.

In summary, for phosphopeptides identified from DDA, TAIMS facilitated phosphosite localization. For unidentified phosphopeptide-spectrum matches near the FDR threshold (1–10% FDR), TAIMS helped salvage hundreds of phosphopeptides with precisely localized phosphosites. Compared to DDA, the DDA-TAIMS workflow identified 21.5% more phosphopeptides (from 5200 to 6320). Under the stricter criterion of localization probability ≥50% and pFind3 PSM score >12, DDA-TAIMS identified 66.6% more high-confidence phosphopeptides than DDA (from 1537 to 2561). When a more stringent filter (localization probability ≥75% and pFind3 PSM score >12) was applied, the improvement was 39.7% (from 1301 to 1817) (Supplementary Figure 16).

## Discussion

In TAIMS, MS2 scans are triggered conditionally when a target precursor ion meets an intensity threshold. TAIMS has a capacity of 2000–4000 targets per run. In contrast, PRM acquires MS2 scans unconditionally following a fixed schedule. PRM has the best chance to capture low-abundance targets if everything else is the same as TAIMS and if the total number of targets do not exceed ∼100 per run.

For a successful implementation of TAIMS, we would like to emphasize the following. First, TAIMS needs reproducible chromatographic retention time, ideally ≤0.5 minutes drift. This is manageable when using the same column and gradient with DDA. To generate the inclusion list, it is recommended to monitor target precursors within a window between ±1 min and ±3 min around the expected elution time depending on the gradient length. Second, dynamic exclusion times are optimized to be 1/7–1/5 of the average peak width, balancing the need for identification quantity and quality (Supplementary Figure 1H).

Our analysis reveals that a significant portion of precursor ions extracted from unidentified DDA CSMs corresponds to "imagined precursors," as their assigned m/z and charge state lack reliable MS1 evidence. This suggests that how MS data are preprocessed can affect profoundly the identification rate of TAIMS data, not only prior to data acquisition (inclusion list) but also after, at the data analysis step. If the isolation-center m/z (the MS2 isolation-window center recorded in the scan header) is assigned as the precursor m/z, as TargetWizard does, only about 10% of targeted precursors yield identification (i.e., at least one MS2 scan results in identification) (Figure 5A). If TAIMS data are preprocessed using pXtract, which tries to locate the monoisotopic peak if it finds the peak at the isolation center is not, the identification rate is increased to 43% (Supplementary Figure 17A). pParse performs the same task, but unlike pXtract, which outputs one monoisotopic peak per MS2, it considers co-eluting ions and may output multiple monoisotopic peaks per MS2. If TAIMS data are preprocessed using pParse, the identification rate rises to approximately 75% (Supplementary Figure 17B). By preprocessing using TargetWizard, we made sure that it was the inclusion targets that were analyzed, not other ions co-isolated with the intended ones.

Although pXtract and pParse led to higher identification rates for precursor ions extracted from unidentified DDA CSMs, the addition of new cross-links remains modest. Specifically, 41 new cross-links were identified, and this number increased to 73 and 88 with pXtract and pParse, respectively—representing less than 7% of the total (1299 cross-links, Supplementary Figure 17). The increase of new cross-link identifications is moderate, because the majority of unidentified CSMs contributing to the inclusion list in Figure 5A correspond to genuine precursor ions already identified in DDA, which are either cross-linked or linear peptides. Therefore, TAIMS following DDA serves to validate DDA identifications, and secondarily to expand cross-link discovery. This highlights the importance of reducing redundancy when generating the inclusion list, a feature that has been incorporated into TargetWizard.

Although in this study, target ions of TAIMS came from DDA analysis, they could originate from alternative sources. For example, an inclusion list of target ions could be generated following DIA^65^ to validate the initial identifications of peptides, PTMs, or cross-links, etc. Additionally, an inclusion list of TAIMS may be created through prediction, such as hypothetical phosphopeptides stemming from biological hypotheses, cross-links predicted from in silico models of protein complexes, or neoantigens predicted from mutanome data (e.g., neudiscMS^66^). We believe that DIA-TAIMS and prediction-TAIMS will open new avenues for MS-based clinical proteomics and are thus worthy of further exploration.

## Methods

### Chemicals

Acetonitrile, formic acid, ammonium bicarbonate and ammonium acetate were purchased from J.T. Baker. Dimethylsulfoxide (DMSO), HEPES, urea, and other general chemicals were purchased from Sigma-Aldrich (St. Louis, MO). Trypsin was purchased from Promega (Wisconsin, WI). Disuccinimidyl sulfoxide (DSSO) was from Pierce.

### Sample preparation

Analysis in this study was primarily focused on DSSO cross-linked *E. coli* samples, with a small portion of the analysis being performed on DSSO cross-linked yeast ribosome samples.

Preparation of *E. coli* lysates: *E. coli* (MG1655) cells were grown at 37°C in 500 mL M9 minimal medium from a 1 mL overnight culture. Cells were harvested after 26 hours at OD600 1.8. Cell lysates were prepared in 50 mM HEPES pH 8.2, 150 mM NaCl using a FastPrep system (MP Biomedicals, Santa Ana, CA) using two volumes of glass beads at 6.5 m/s, 20 s per pulse for four pulses, with 5 minutes of cooling on ice between pulses. The lysates were cleared by centrifugation at top speed in a tabletop microfuge for 30 min. Protein concentrations were determined using the bicinchoninic acid assay.

Preparation of yeast ribosome extracts: Yeast cultures were initially grown on YPD. The overnight culture was diluted to an OD600 of 0.15 and further grown in 1 L at 30 until an OD600 of 1.0 was reached. Cells were then collected by centrifugation at 4000 g for 15 minutes at 4 and washed with 100 ml of pre-cooled lysis buffer (30 mM Tris-acetate, pH 7.4, 100 mM NH_4_Cl, 12 mM MgCl_2_, 2 mM DTT, 1% PMSF). The washed cells were resuspended in lysis buffer containing protease inhibitor, disrupted by a high-pressure homogenizer three times, and centrifuged at 14,000 rpm for 1 hour at 4 to obtain the supernatant. This supernatant was layered onto a sucrose cushion (1 M sucrose, 50 mM Tris-acetate, pH 7.4, 500 mM NH_4_Cl, 12 mM MgCl_2_, 2 mM DTT) and centrifuged at 30,000 rpm for 20 hours at 4 □. The resulting precipitate was resuspended in lysis buffer to obtain crude ribosomes. These ribosomes were loaded onto a linear 10-50% sucrose density gardient and centrifuged at 28,700 rpm for 7 hours at 4 □. Fractions were collected based on UV260 absorbance peaks detected using a Nanodrop spectrophotometer, concentrated using ultrafiltration tubes with displacement buffer, and stored at -80 □ after determining protein concentration using a BCA assay.

Preparation of phosphorylated sample: The mice brain tissues were homogenized in 1% SDS lysis buffer supplemented with protease inhibitor (4693132001, Merck) and phosphatase inhibitor (4906837001, Merck). The tissues were further dissociated by a cryogenic tissue grinder (JXFSTPRP-CLN-24, Shanghai Jingxin). The cell suspension was collected by centrifuge at 12,000 g for 10 minutes at 4□. The protein mixtures were then reduced with 100 mM dithiothreitol (DTT, Roche) at 56□ for 30 minutes and followed by alkylation in 100 mM iodoacetamide (IAA, TCI) in the dark for 1 h. Then proteins were precipitated with four times the sample volume of cold (-20□) acetone overnight. The protein pellet was collected by centrifuge at 12,000 g for 10 minutes at 4□. The protein pellet was resuspended with 8 M urea. After that, the protein mixtures were digested into peptides with trypsin using the filter-aided sample preparation (FASP) method. The peptides were desalted with tC18 SepPak columns (100 mg, Waters), and the eluents were vacuum-dried. Phosphopeptide enrichment was performed with a Fe-NTA phosphopeptide enrichment kit (Thermo Scientific) following the manufacturer’s protocol.

### Protein cross-linking

*E. coli* cell lysates or yeast ribosome extracts were adjusted to a protein concentration of 1 mg/mL and incubated with 1 mM DSSO for 45 minutes at room temperature. The cross-linking reaction was then stopped by adding 20 mM ammonium bicarbonate. Next, the proteins were precipitated by adding ice-cold acetone, followed by air drying. Finally, the proteins were resuspended in a solution containing 8 M urea and 100 mM Tris at pH 8.5 and subsequently digested with trypsin. Digested peptides were desalted using peptide desalting columns.

### Liquid Chromatography

Cross-linked samples were analyzed with an EASY-nLC 1200 system (Thermo Fisher Scientific, Waltham, MA) interfaced with a Fusion Lumos mass spectrometer (Thermo Fisher Scientific). A one-column setup with an analytical column (150 μm, 35 cm, C18, 1.9 μm) of a 5 μm tip was applied. For DSSO cross-linked *E. coli* samples, peptides were eluted with a gradient from 2% to 6% buffer B in 4 min, then to 40% B in 156 min, and finally to 100% B in 10 min, held for 5 min, and then back to 10% B in 5 min, at a constant flow rate of 450 nL/min. For DSSO cross-linked yeast ribosome samples, the elution gradient was adjusted for a shorter LC/MS acquisition: 2-6% B in 4 min, 6-40% B in 96 min, 40-100% B in 10 min, held at 100% B for 5 min, and then back to 10% B in 5 min, at a constant flow rate of 450 nL/min.

Phosphorylated sample was analyzed with an EASY-nLC 1000 system (Thermo Fisher Scientific, Waltham, MA) interfaced with a Q Exactive HF mass spectrometer (Thermo Fisher Scientific). An analytical column (150 μm, 50 cm, 1.9 μm C18) of a 5 μm tip was applied. Peptides were eluted with the following gradient: 2 to 6% buffer B in 3 min, 6 to 28% B in 55 min, 28 to 80% B in 10 min, 80% in 7 min, at a constant flow rate of 600 nL/min.

### DDA Mass Spectrometry

For cross-linked samples: a Fusion Lumos mass spectrometer (Thermo Fisher Scientific) was utilized. MS1 scan resolution was set to 120,000 with an AGC (automatic gain control) target of 1 × 10^6^ and a maximum ion injection time of 100 ms. The scan range (m/z) was set from 400 to 1600. The most abundant precursor ions with z = 3–6, passing the peptide match filter ("monoisotopic precursor selection"), were selected for MS2 within a cycle time of 2 s. MS2 scan resolution was set to 30,000 with an AGC target of 1 × 10^5^ and a maximum ion injection time of 120 ms. The quadrupole isolation window was set to 1.6 m/z, and the normalized collision energy of stepped HCD was set at 27 ± 6. Dynamic exclusion was enabled for 45 s after a single count and included isotopes. LC-MS acquisition lasted 180 minutes for *E. coli* sample and 120 minutes for yeast ribosome.

For the phosphorylated sample: a Q Exactive HF mass spectrometer (Thermo Fisher Scientific) was utilized. MS1 scan resolution was set to 120,000 with an AGC (automatic gain control) target of 3 × 10^6^ and a maximum ion injection time of 50 ms. The scan range (m/z) was set from 400 to 1600. The most abundant 15 precursor ions with z = 2–5 were selected for MS2 analysis. MS2 scan resolution was set to 15,000 with an AGC target of 1 × 10^5^ and a maximum ion injection time of 60 ms. The quadrupole isolation window was set to 1.6 m/z, and the normalized collision energy of HCD was set at 27. Dynamic exclusion was enabled for 45 s after a single count and included isotopes. Each LC-MS acquisition lasted 75 minutes.

### Inclusion list generation

Inclusion lists were created with TargetWizard, specifying the monoisotopic m/z, charge state, and monitoring time window for each target precursor. The monitoring window was centered around the DDA retention time, which corresponds to the apex of the precursor’s extracted ion chromatogram (XIC).

For TAIMS-250 and TAIMS-500 on DSSO cross-linked *E. coli* lysates, the monitoring window was ±3 minutes. An individualized intensity threshold of 10% was set to trigger MS2 scans; specifically, an MS2 scan was initiated whenever the precursor’s intensity exceeded 10% of its peak intensity previously recorded in DDA mode.

For DSSO cross-linked yeast ribosome samples, the retention time windows were set to ±2 minutes.

For phosphorylated sample, among the phosphorylated peptides identified by pFind3 under a 1% peptide FDR, there were 1567 peptides with a PTMiner site localization probability falling below the reliability threshold of 0.5. In addition, 637 phosphorylated peptides have a peptide FDR ranging from 1% to 10%. From both these subsets of peptides, we selectively chose the best precursor ions based on the raw score and compiled them into an inclusion list. Monitoring time windows of ±2 minutes around the DDA retention time of the optimal precursor were applied.

### TAIMS Mass Spectrometry

a Fusion Lumos mass spectrometer (Thermo Fisher Scientific) was used. MS1 scan resolution was set to 120,000 with an AGC (automatic gain control) target of 1 × 10^6^ and a maximum ion injection time of 100 ms. The scan range (m/z) was set from 400 to 1600. The most abundant precursor ions on the inclusion list, meeting the retention time requirements, were selected for MS2 using Top Speed acquisition mode with a cycle time of 2 s. The peptide match filter ("monoisotopic precursor selection") was inactivated. MS2 scan resolution was set to 120,000 with an AGC target of 2 × 10^5^ and a maximum ion injection time of 246 ms on TAIMS-250 and 502 ms on TAIMS-500. The quadrupole isolation window was set to 1.6 m/z, and the normalized collision energy of stepped HCD was set at 27 ± 6. Dynamic exclusion was enabled for 3 s after a single count and included isotopes. Each LC-MS acquisition lasted 180 minutes.

For phosphorylated sample: a Q Exactive HF mass spectrometer (Thermo Fisher Scientific) was utilized. The Mass spectrometric parameters used were as follows: MS1 scan resolution was set to 120,000 with an AGC target of 3 × 10^6^ and a maximum ion injection time of 50 ms. The scan range (m/z) was set from 400 to 1600. The most abundant 15 precursor ions on the inclusion list, meeting the retention time requirements, were selected for MS2 analysis. MS2 scan resolution was set to 120,000 with an AGC target of 2 × 10^5^ and a maximum ion injection time of 246 ms. The quadrupole isolation window was set to 1.6 m/z, and the normalized collision energy of HCD was set at 27. Dynamic exclusion was enabled for 3 s after a single count and included isotopes. Each LC-MS acquisition lasted 75 minutes.

### Identification of cross-linked peptides

For preprocessing mass spectrometry raw files, pParse was utilized for DDA data, while TargetWizard was primarily employed for TAIMS data. Additionally, pXtract and pParse were tested with TAIMS data for evaluation. For each MS2 scan in DDA or TAIMS, pParse outputs the monoisotopic peak information (including m/z and charge state) for all software-calibrated precursors, including the MS2 isolation window center-associated precursor and any co-eluting precursors relevant to that scan. pXtract provides the instrument-calibrated monoisotopic peak information specifically for the MS2 isolation window center-associated precursor. In contrast, TargetWizard outputs the specified m/z and charge values from the inclusion list as the monoisotopic peak information for the corresponding precursors.

Cross-linked peptides were identified using in-house software pLink 2.4.2, dedicated to DSSO identification (pLink 3.0.17 compatibility was verified; see Supplementary Figure 10 and main text). The search parameters were as follows: Flow Type, gas phase cleavable (stepped HCD); cross-linker, DSSO; peptide length, from 6 to 60 amino acids; digestion, trypsin with up to three missed cleavages; mass tolerance for precursor ions, ±5 ppm for both DDA and TAIMS; mass tolerance for fragment ions, ±20 ppm for DDA and ±5 ppm for TAIMS. For the yeast ribosome DDA data described in the supplementary information, mass tolerance for fragment ions was adjusted to ±10 ppm for cross-link identification, and ±20 ppm was used to calculate FIC and site-FIC.

Other search parameters on *E. coli* data and yeast ribosome data: K, S, T, Y and protein N-termini were set as cross-linkable sites for both peptides; following modifications were considered: Carbamidomethylation of cysteine as a fixed modification and Methionine oxidation as a variable modification. Results were filtered by applying a 5% FDR cutoff at precursor level. FDR was controlled separately for intra-protein and inter-protein cross-links.

### Phosphorylated sites identification and localization

For DDA data, pParse was used to extract monoisotopic m/z from mass spectrometry raw file. For TAIMS data, pXtract was used to extract monoisotopic m/z from mass spectrometry raw file, as it provided higher identification sensitivity for phosphopeptides in our setup. All phosphorylation sites were identified by pFind3 in the restricted search mode. Carbamidomethylation on cysteine was set as a fixed modification; Phosphorylation on S/T/Y and oxidation on methionine were also set as variable modifications. The mouse proteome was used as a database. Trypsin digestion was allowed up to three missed cleavages. Results were filtered using 1% peptide level FDR. For other items, the default settings were retained.

For phosphorylated sites localization: the pFind3 search parameter file “pFind.cfg” and identification file “pFind.spectra” were imported to PTMiner 1.2.7, and posterior probability was used for the localization of modification sites.

### Inclusion list generation and post-TAIMS data analysis using TargetWizard

TargetWizard, a user-friendly open-source tool, streamlines TAIMS analysis and requires no coding skills. Its key functions include:

*1. Inclusion list compilation:* TargetWizard assists in compiling an inclusion list, specifying the precursor ions to be monitored during the TAIMS acquisition.
*2. Comprehensive Post-TAIMS data analysis:* TargetWizard is invaluable for comprehensive post-TAIMS data analysis. Specifically, for each target precursor ion listed for inclusion, the software finds all MS2 spectra that align with the m/z, charge, and retention time of the target precursor ion. Additionally, it calculates the similarity between the current TAIMS spectra and the best DDA spectra of the target precursor ion. This tool summarizes identification results for all these spectra.
*3. Fragment-ion-coverage & Signal-to-Noise output:* TargetWizard computes fragment-ion-coverage and extracts signal-to-noise data for the identified cross-linked spectra, thereby offering a solid analytical foundation for TAIMS experiments.

Detailed information on TargetWizard’s functions, as well as instructions on how to download the software, is available here: https://targetwizard.ctarn.io.

### Accurate coupling of target ions with TAIMS MS2 spectra

TargetWizard first associates TAIMS-acquired MS2 spectra with inclusion list targets based on m/z, charge, and retention time. To verify that the isolated precursor in the TAIMS MS2 spectrum matches the intended target, TargetWizard calculates a similarity score. For each target precursor ion, fragment ion peaks matched to the corresponding DDA spectrum are extracted as reference peaks. The fraction of these reference peaks detected in the matched TAIMS MS2 spectrum serves as the similarity measure.

TAIMS MS2 spectra with similarity exceeding a defined threshold are considered to represent the fragmentation spectra of the intended precursor derived from DDA.

The similarity threshold is automatically determined per dataset using carefully constructed positive and negative samples. Positive samples include DDA–TAIMS precursor pairs aligned within 10 ppm m/z and a 5-minute retention time window, with matching high-confidence MS2 identifications (high-FIC). Negative samples pair the same DDA precursor with TAIMS spectra randomly chosen from retention times at least ±15 minutes apart to ensure no shared origin.

Method performance was evaluated by false positive rate (FPR) and false negative rate (FNR), representing incorrect assignments and missed TAIMS hits, respectively. The threshold was set to achieve a 1% FNR, with an observed FPR below 1%, ensuring reliable coupling between target precursor ions and their corresponding MS2 spectra.

## Acknowledgements

The authors gratefully acknowledge Dr. Yue Zhao for providing phosphorylated samples from mouse brain and Dr. Jian-Hua Wang for providing yeast ribosome extracts. This work was supported by the National Natural Science Foundation of China (32071435 to S.-M.H.), the National Key Research and Development Program of China (Grant No. 2020YFF01014505), and the intramural grants of NIBS, Beijing, which is funded by the Ministry of Science and Technology of China, the municipal government of Beijing, and Tsinghua University.

## Author contributions

A.M. optimized the mass spectrometry method, analyzed and interpreted the data, and wrote the manuscript. C.T. developed TargetWizard. M.-Q.D. guided the study, interpreted the data, and wrote the manuscript. S.-M.H. conceived the idea of improving MS2 quality using targeted MS, guided the study, interpreted the data, and revised the manuscript. P.-Z.M. developed a Python tool for accurate coupling of target ions with TAIMS MS2 spectra. Z.-L.C. and P.-Z.M. developed pLink 2.4.2. H.C. and Y.C. helped with experimental design.

## Data availability

All datasets generated and analyzed in this study are summarized in Supplementary Table 2 and have been deposited to the ProteomeXchange Consortium (https://proteomecentral.proteomexchange.org) via the iProX partner repository under the dataset identifier PXD065604. Reviewers can access the data through the following link using the password provided: https://www.iprox.cn/page/PSV023.html;?url=1774992819586xPpy (password: D62J)

## Code availability

The pLink 2.4.2 software used in this study is accessible via ProteomeXchange under the project identifier PXD065604. The Python tool developed to determine the similarity threshold for calculating the target triggering rate is available at: https://github.com/NorthblueM/TAIMS-paper-scripts. Additionally, the software and source code for TargetWizard can be found at http://targetwizard.ctarn.io.

