## Supporting Information for "Improved identification of peptides, modification sites, and cross-link sites by Target-enhanced Accurate Inclusion Mass Screening (TAIMS)"

### Index to Supplementary Figures and Tables

**Supplementary Figure 1.** Optimization of MS parameters to improve spectral quality. 3

**Supplementary Figure 2.** Effect of fragment ion mass tolerance on the quality of cross-link identifications. 5

**Supplementary Figure 3.**"Enrichment" of target precursors at the MS2 level. 6

**Supplementary Figure 4.** TAIMS improves spectral quality of DSS cross-links. 7

**Supplementary Figure 5.** Distribution of fragment ion coverage of true vs false CSMs. 8

**Supplementary Figure 6.** A false DDA identification pinpointed by TAIMS. 9

**Supplementary Figure 7.** Performance of TAIMS on a DSSO cross-linked yeast ribosome sample. 10

**Supplementary Figure 8.** Schematic explanation of the overlap between the DDA identifications and the TAIMS identifications. 11

**Supplementary Figure 9.** An inter-protein cross-link identification salvaged by TAIMS. 13

**Supplementary Figure 10**. Consistent increase in high-FIC cross-link identifications by TAIMS with pLink 2 and pLink 3. 14

**Supplementary Figure 11.** An example of TAIMS spectra resisting sensitivity loss in cross-link identification caused by search space expansion. 16

**Supplementary Figure 12.** Statistics showing resistance of TAIMS spectra to sensitivity loss from search space expansion in a DSSO cross-linked yeast ribosome sample. 17

**Supplementary Figure 13.** Prioritization of candidate targets for TAIMS analysis. 18

**Supplementary Figure 14.** Multi-batch scheduling in TargetWizard. 20

**Supplementary Figure 15.** TAIMS improves the localization accuracy of cross-linked sites. 21

**Supplementary Figure 16.** Phosphosite localization using a probability threshold of 75%. 22

**Supplementary Figure 17.** TAIMS analysis outcome of subthreshold DDA entries using preprocessing tools alternative to TargetWizard. 23

**Supplementary Table 1.** Recommendation on how to shorten an inclusion list for TAIMS 24

**Supplementary Table 2.** Data analyses and datasets used in this study 25

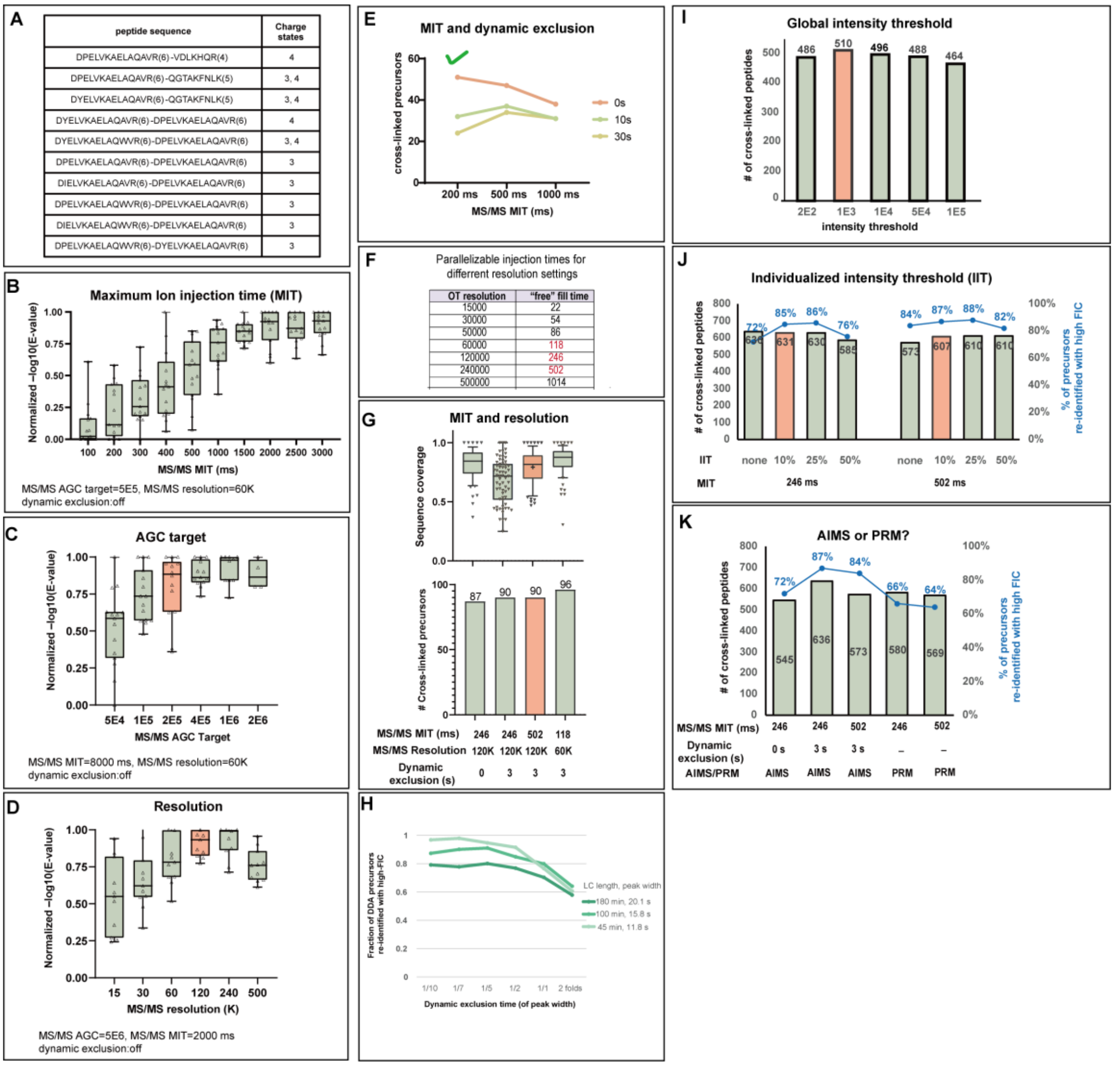


*Supplementary Figure 1. Optimization of MS parameters to improve spectral quality.*

**(A)** List of the chemically synthesized cross-linked peptide precursor ions used in this analysis. **(B–D)** Effects of maximum injection time (MIT) (B), AGC target (C), and resolution (D) on MS2 spectral quality. Each data point corresponds to an individual cross-linked peptide precursor. Spectral quality is represented as the –Log10(E-value) normalized to the maximum value within each experimental group. **(E)** Effect of dynamic exclusion time on the number of cross-link identifications. DDA analysis was initially performed on a BSA sample cross-linked with DSSO. The resulting raw file was searched against the sequence of BSA without decoy sequences, and a total of 497 cross-link precursors were "identified" without FDR control. From these precursors, less reliable ones with an *E*-value>1E-5 were compiled into an inclusion list for AIMS analysis. Nine separate AIMS experiments were conducted, each with different MIT/dynamic exclusion settings but utilizing the same inclusion list. The resulting AIMS raw files were then searched against the sequences of BSA and 293 common contaminant proteins for cross-link identification, applying a 5% FDR threshold for CSMs. **(F)** Parallelization of MIT and resolution on a Fusion Lumos Mass Spectrometer. Available resolution settings for the Fusion Lumos mass spectrometer and the corresponding maximum ion injection times that can be employed without wasting any orbitrap time. Injection time setting examined are colored red. **(G)** Optimization of MIT for MS2. The DDA data of a DSSO treated BSA sample was searched against sequences of BSA and 293 common contaminant proteins with a 5% FDR at the CSM level. The inclusion list for subsequent TAIMS analysis contained 88 cross-link precursors that were identified with low confidence (E-value>1E-8 or peptide bond cleavage<0.6 for either of the alpha or beta peptide) and 232 precursors that were filtered out by 5% FDR threshold. Four AIMS experiments were conducted with varied MIT and resolution settings. The resulting AIMS data were searched in the same way as the DDA data. **(H)** Effect of dynamic exclusion time on TAIMS performance across different gradient lengths. The inclusion list of AIMS consisted of the precursor ions of CSMs, all of which come from a pLink search of the DDA data of a DSSO cross-linked *E. coli* lysate. They were exclusively forward database matches, but without FDR control. TAIMS experiments were conducted with dynamic exclusion times set to 1/10–2-fold of the mean peak width for each gradient length. Peak widths were calculated using Skyline 21.2.0 based on identified cross-linked peptide pairs (under 5% FDR) from DDA. The mean peak widths for gradient lengths of 180 min, 100 min, and 45 min were 20.1 s, 15.8 s, and 11.8 s, respectively. Targeted MS performance was assessed based on the total number of identified cross-link precursors and the high-FIC re-identification rate of cross-link precursors initially identified by DDA. **(I)** Optimization of global intensity threshold. The sample and the inclusion list are the same as in Supplementary Figure 1H. The performance of targeted MS was evaluated based on the total number of cross-link precursors identified. **(J)** Optimization of individualized intensity threshold (IIT). In AIMS analysis, the intensity threshold to trigger a MS2 scan can be specified for individual precursor ions, expressed as a percentage of a precursor's DDA intensity. The inclusion list consisted of 1384 CSMs generated from a pLink search of the DDA data of a yeast ribosome sample cross-linked using DSSO. They were exclusively forward database matches, but without FDR control. Targeted MS performance was assessed based on the total number of identified cross-link precursors and the high-FIC re-identification rate of cross-link precursors initially identified by DDA. **(K)** Comparison of AIMS and PRM. The sample and the inclusion list are the same as in (J). For panels H, I, J and K, the AIMS or PRM data were searched against the sequences of *E. coli* or yeast ribosome proteins with a 5% FDR at the precursor level.

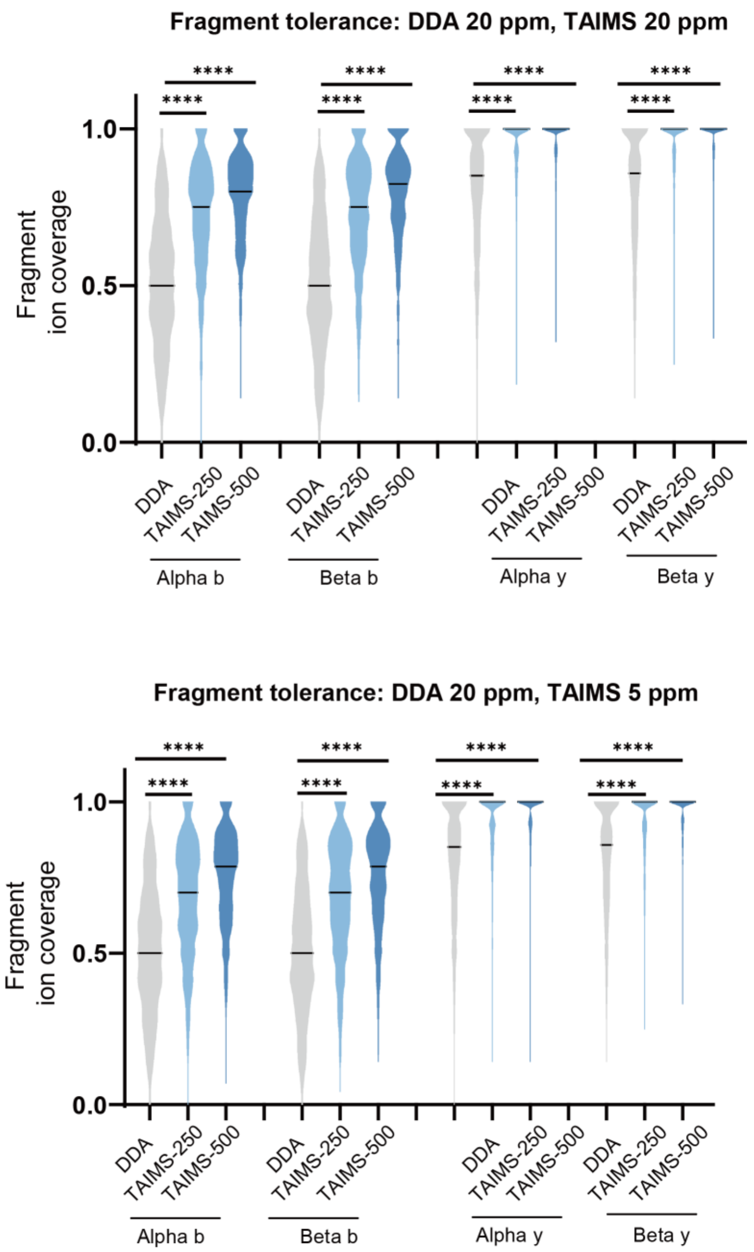


*Supplementary Figure 2. Effect of fragment ion mass tolerance on the quality of cross-link identifications.*

Fragment ion mass tolerance of ±5 ppm or ±20 ppm in post-search analysis made little difference. For pLink2 searches, ±20 ppm fragment ion tolerance was used for DDA and ±5 ppm for TAIMS. To calculate fragment ion coverage of the identified spectra, ±20 ppm was applied for DDA, and either ±20 ppm (upper panel) or ±5 ppm (lower panel) was used for TAIMS.

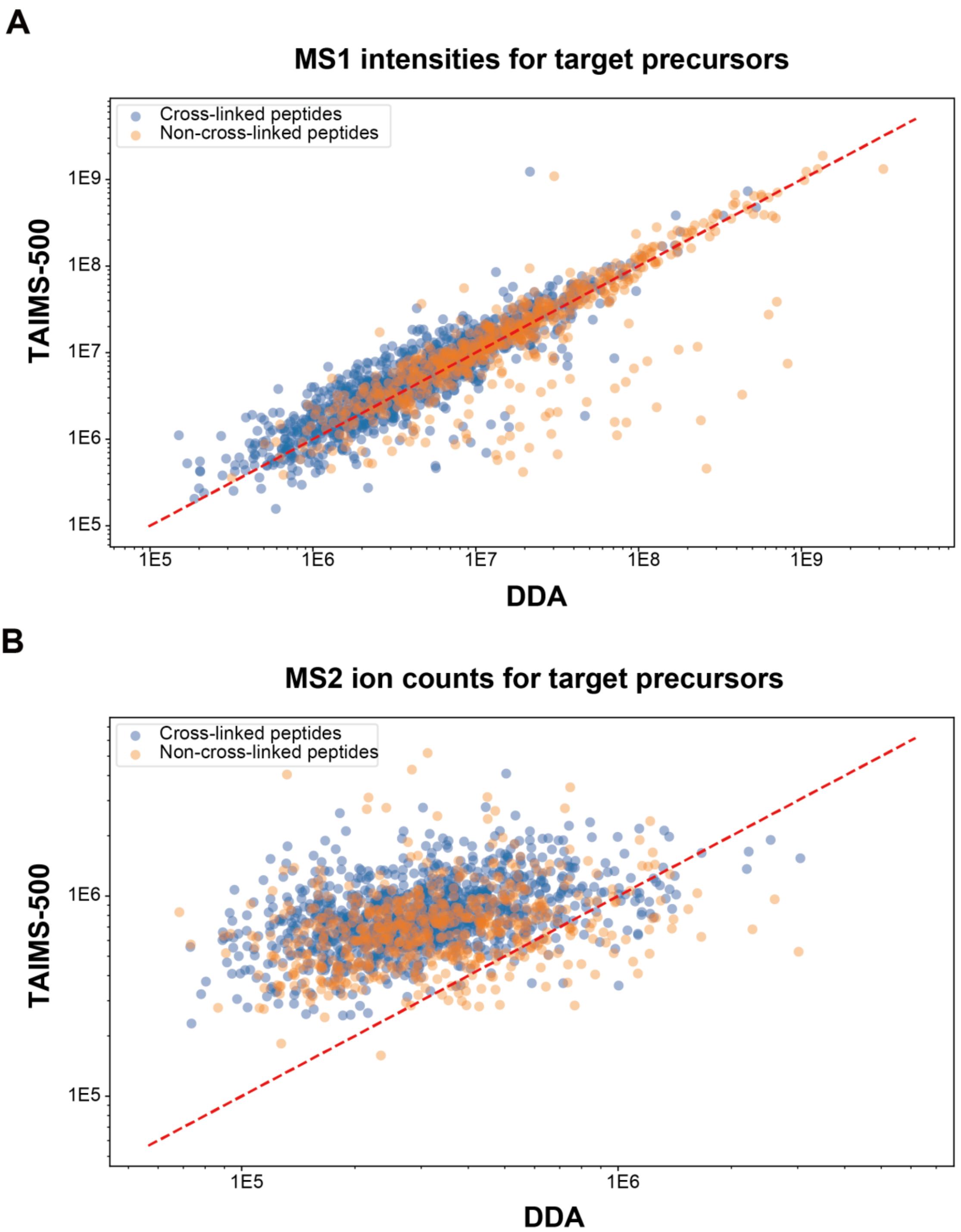


*Supplementary Figure 3.* *"Enrichment" of target precursors at the MS2 level.*

**(A)** Target precursors identified by both DDA and TAIMS-500 have essentially the same intensities in MS1. (**B**) For most of the target precursors shown in (A), their MS2 ion counts are higher in the TAIMS data than in the DDA data.

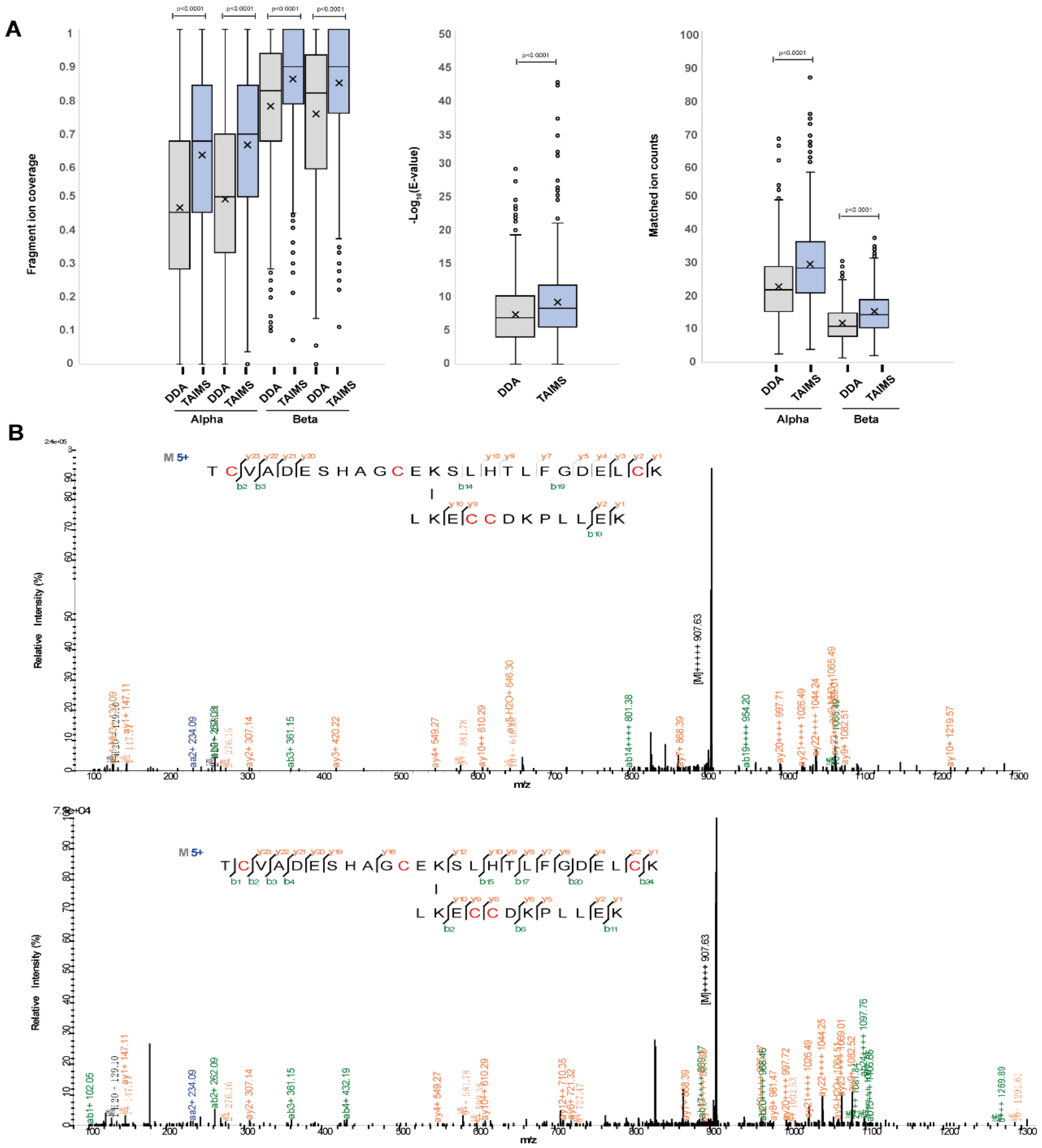


*Supplementary Figure 4. TAIMS improves spectral quality of DSS cross-links.*

A common set of 599 cross-link precursor ions was identified from a DSS-linked BSA sample across two MS runs, one DDA and one TAIMS. To compare among the cognate MS2 spectra, CSMs with the best *E*-values were chosen, one each from DDA and TAIMS. For pLink2 search of the DDA and the TAIMS data, fragment ion mass tolerance was set to ±20 ppm and ±5 ppm, respectively. **(A)** Comparison of 599 sets of cognate DDA and TAIMS MS2 spectra with respect to the indicated spectral quality metric. To calculate fragment ion coverage for both DDA and TAIMS spectra, mass tolerance of ±20 ppm was used. Statistical significance was assessed using the Wilcoxon matched-pairs test. **(B)** Example of MS2 quality improvement in TAIMS. A mass tolerance of 20 ppm was used for annotation.

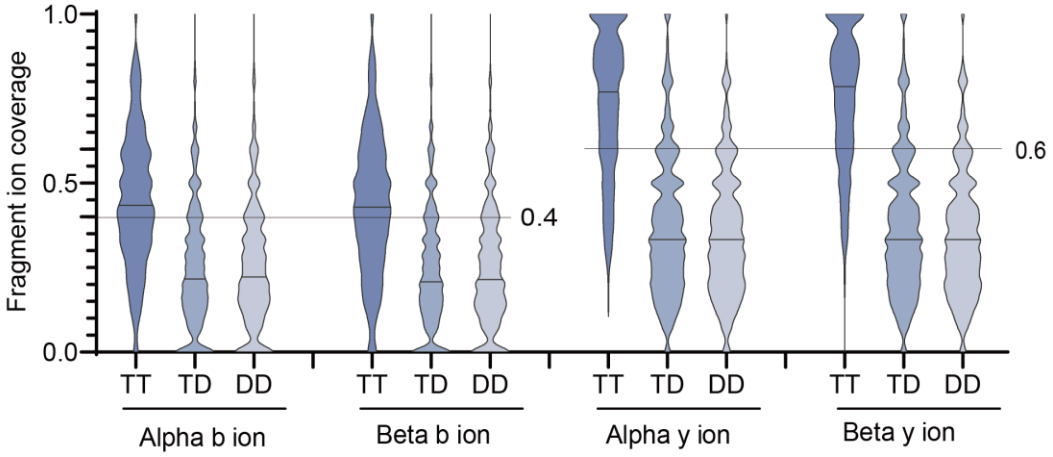


*Supplementary Figure 5. Distribution of fragment ion coverage of true vs false CSMs.*

The *E. coli* DDA data was analyzed to find how to separate reliable cross-link identifications from false ones according to b/y ion fragment coverage. TT, CSMs with α- and β-peptides both in the target sequence database and with *q*-value of zero (highly reliable); TD and DD, CSMs with one or both peptides in the decoy sequence database. Fragment ion mass tolerance is ±20 ppm.

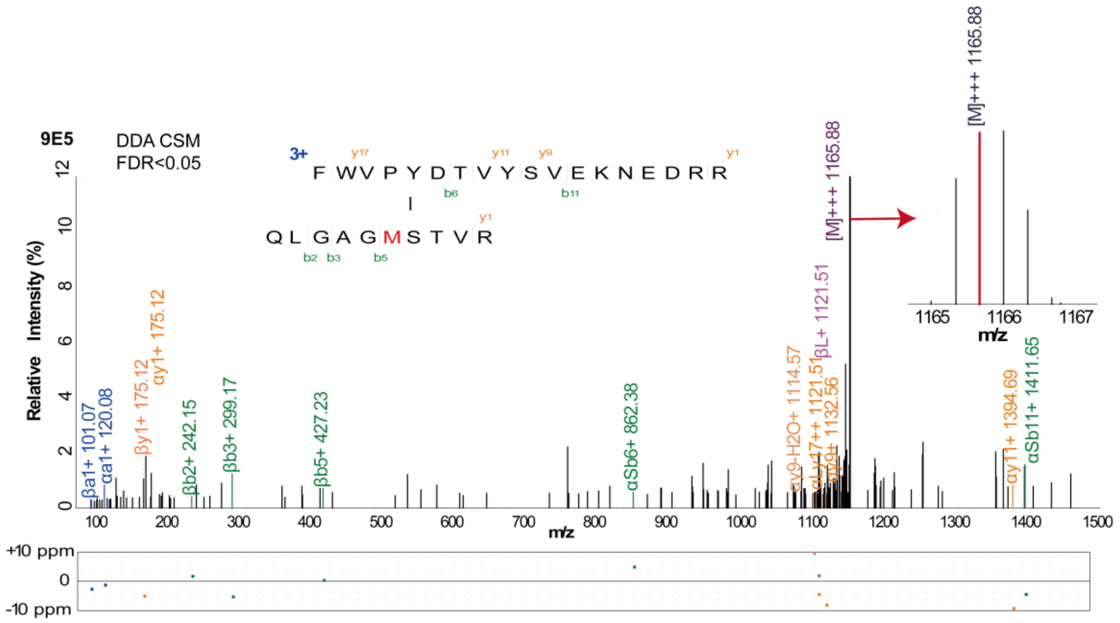


*Supplementary Figure 6.* A false DDA identification pinpointed by TAIMS.

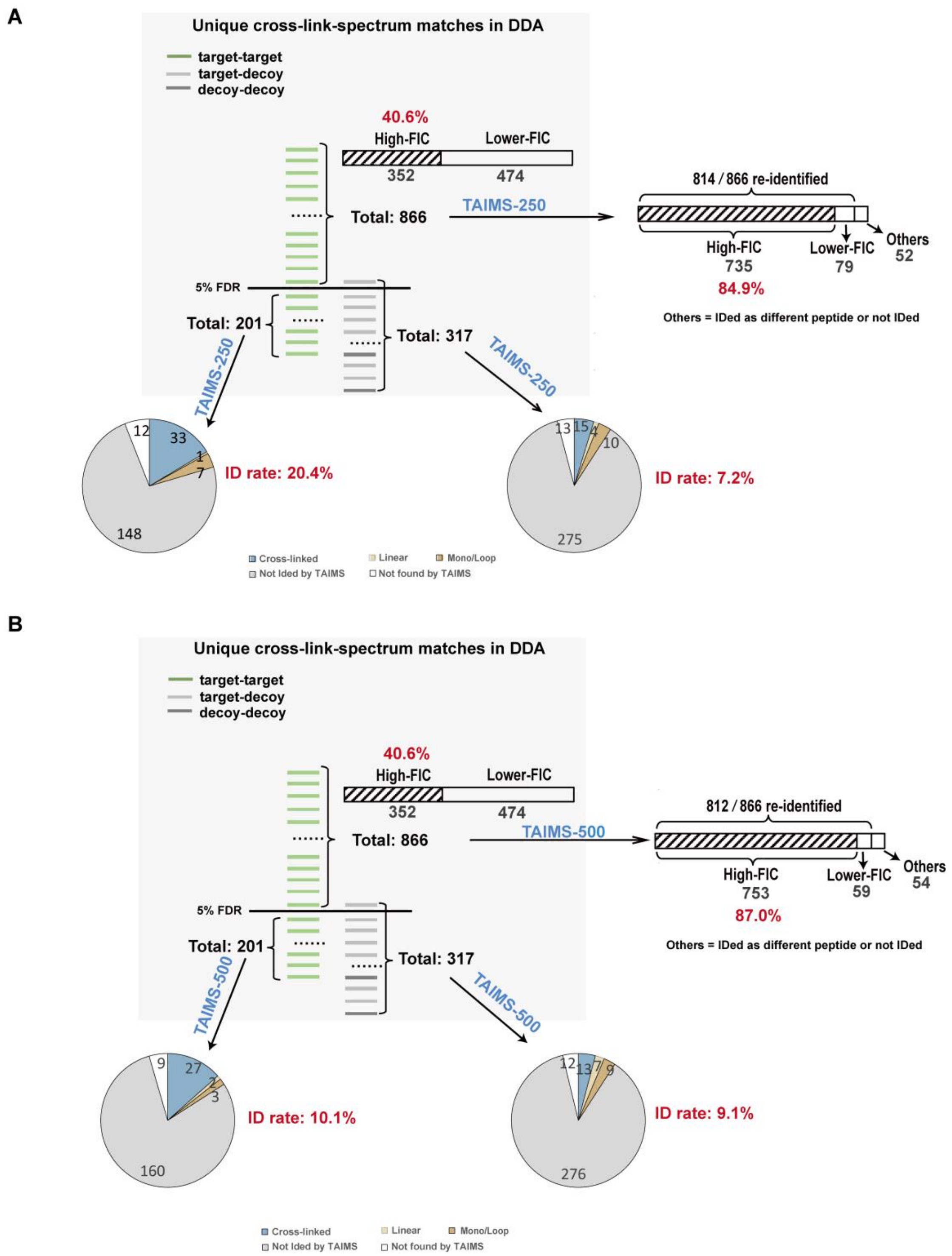


*Supplementary Figure 7. Performance of TAIMS on a DSSO cross-linked yeast ribosome sample.* Initial DDA analysis revealed 1384 unique CSMs via pLink2. Precursor ions from these CSMs comprised the inclusion list for subsequent TAIMS acquisition. (**A**) and (**B**) display the outcome of TAIMS-250 and TAIMS-500 analysis, respectively. Data were preprocessed with TargetWizard. A mass tolerance of ±20 ppm was used for FIC calculation.

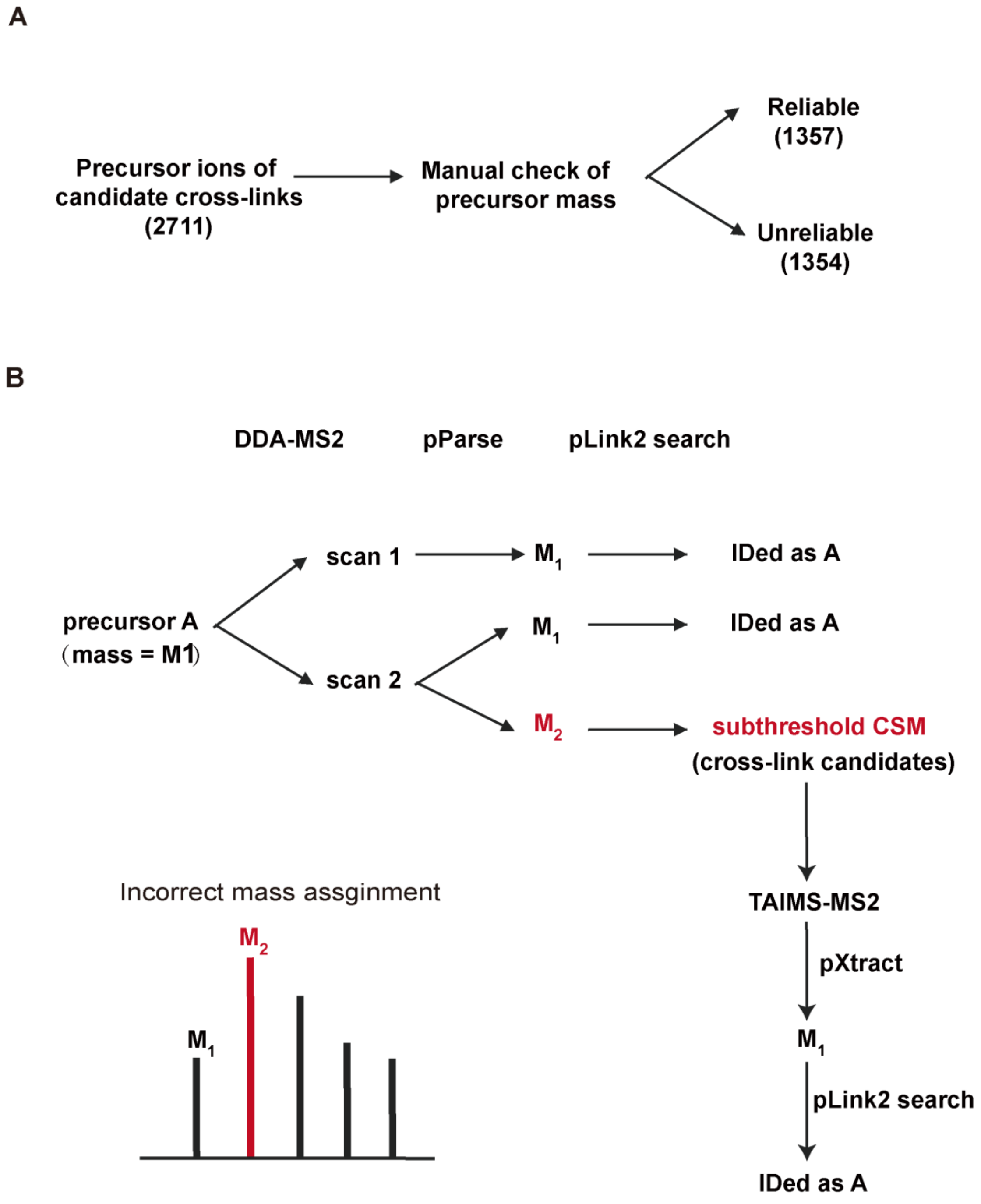


*Supplementary Figure 8. Schematic explanation of the overlap between the DDA identifications and the TAIMS identifications.*

**(A)** The precursor ion masses of 2711 candidate CSMs from the inclusion list of TAIMS-250 analysis were manually verified using their MS1 spectra. Based on this verification, they were classified as either reliable or unreliable assignments. Unreliable precursor masses arose from errors in precursor assignment during the initial DDA-MS2 data analysis. In our dataset, misassignments were caused by either erroneous monoisotopic peak determination (search engines typically test multiple isotopic masses for each MS2 scan to enhance identification sensitivity) or incorrect charge state determination. **(B)** Illustration of the mechanism underlying the overlap between TAIMS identifications from the candidate pool and initial DDA identifications. Precursor A is analyzed in two MS2 scans: it is correctly identified in Scan 1. In Scan 2, two precursor masses are extracted as monoisotopic masses: $M_1$ (genuine precursor) and $M_2$ (imagined precursor). The latter generates a subthreshold candidate CSM that is subsequently included in the inclusion list. During TAIMS, because $M_1$ and $M_2$ share the same isolation window, MS2 scan triggered by $M_2$ leads to fragmentation of $M_1$. If TargetWizard, pXtract, and pParse are used to preprocess the resulting MS2 spectrum, the pLink identification outcome is expected to be none, Precursor A, or Precursor A and possibly Precursor B, respectively. As such, there is a substantial overlap between the identifications derived from these targets and the initial DDA data.

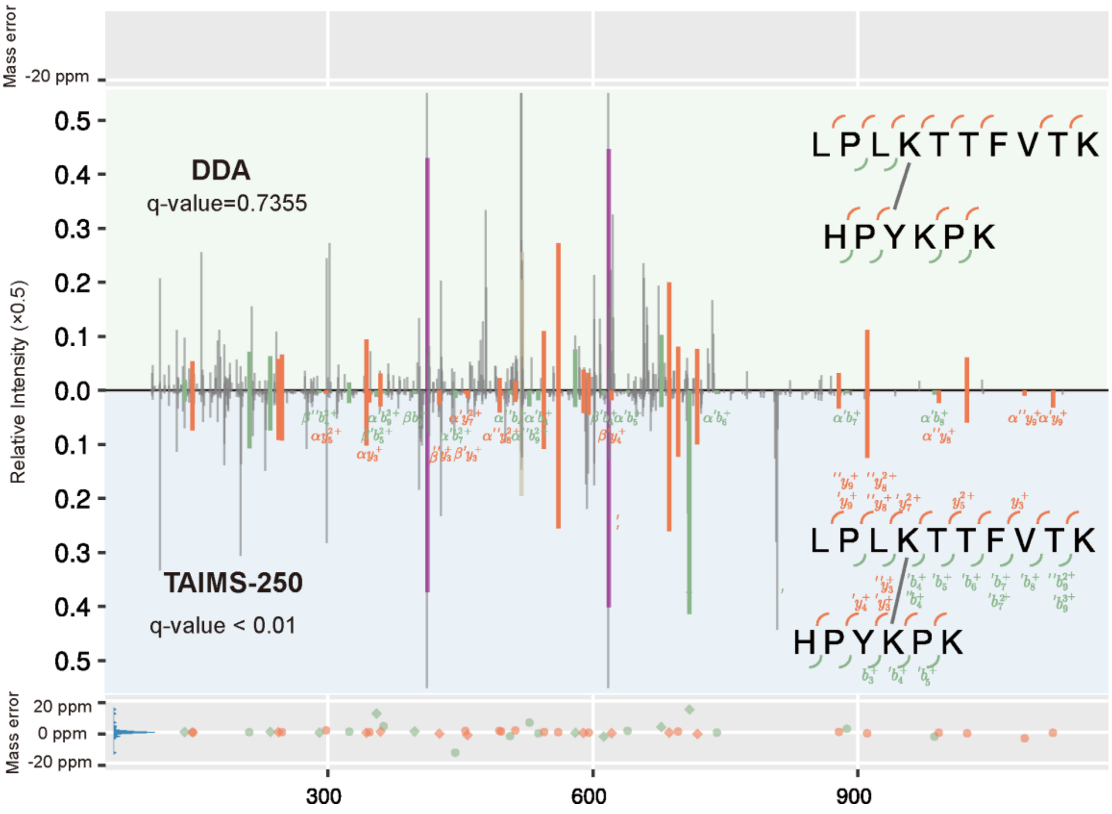


*Supplementary Figure 9.* *An inter-protein cross-link identification salvaged by TAIMS.*

The best DDA CSM contrasted against the best TAIMS CSM of the same precursor ion. Of the fragment ions, only those uniquely detected in the DDA MS2 or the TAIMS MS2 are labeled.

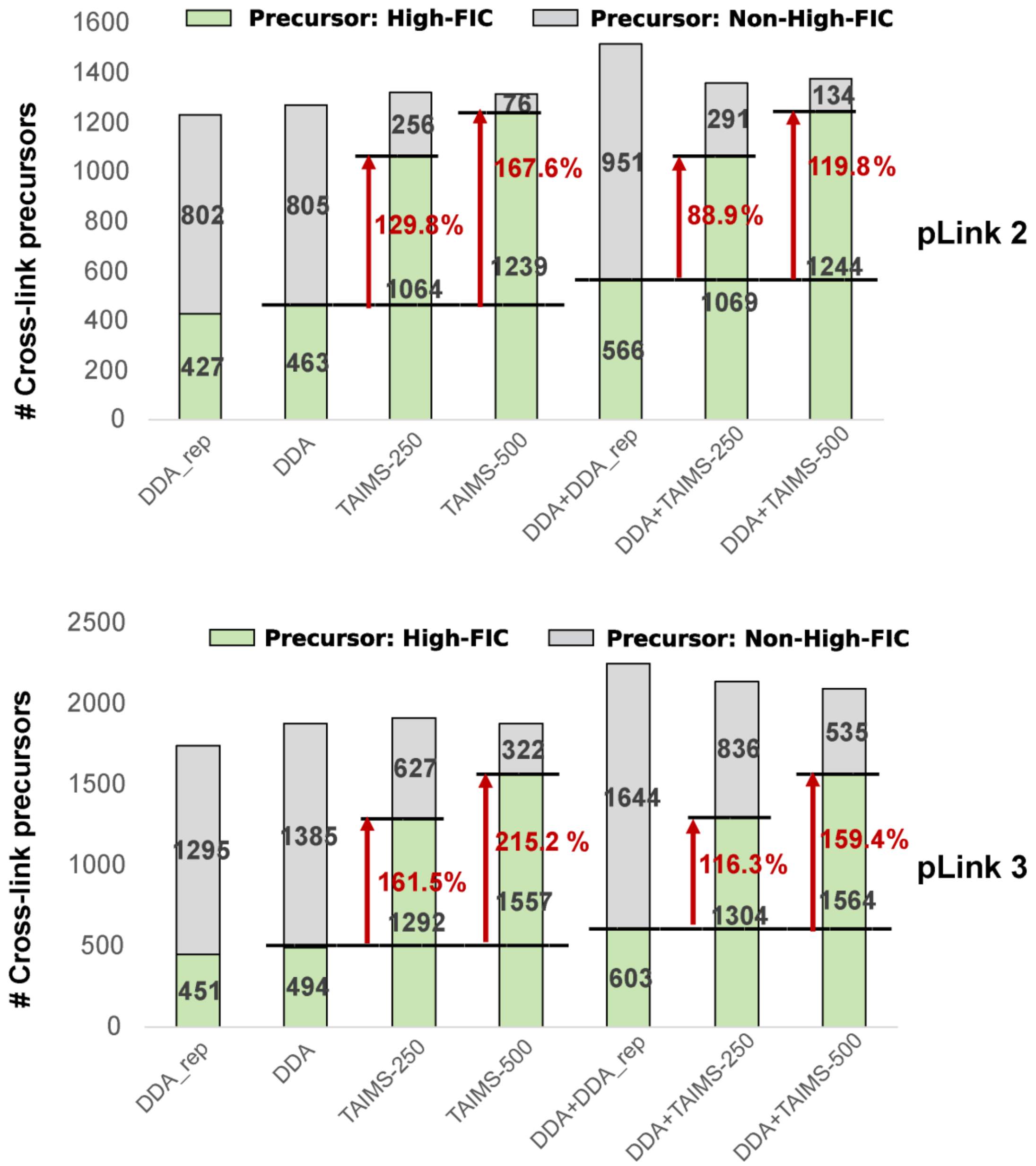


*Supplementary Figure 10. Consistent increase in high-FIC cross-link identifications by TAIMS with pLink 2 and pLink 3.*

To assess whether the benefits of TAIMS are specific to pLink 2.4.2, we performed a parallel DDA–TAIMS experiment on the DSSO-crosslinked yeast ribosome sample using pLink 3.0.17. DDA data were searched separately with pLink 2.4.2 and pLink 3.0.17 at 100% precursor FDR, and inclusion lists were generated from each search using TargetWizard. The pLink 2 search yielded 1654 cross-link precursors; the pLink 3 search yielded 31,907, from which only FDR < 20% precursors (2404) were retained for the inclusion list to enable a side-by-side comparison. TAIMS data were acquired using both inclusion lists and searched with pLink 3.0.17 at 5% precursor FDR. Results are shown before and after applying the high-FIC requirement. FIC values were calculated with a mass tolerance of ±20 ppm.

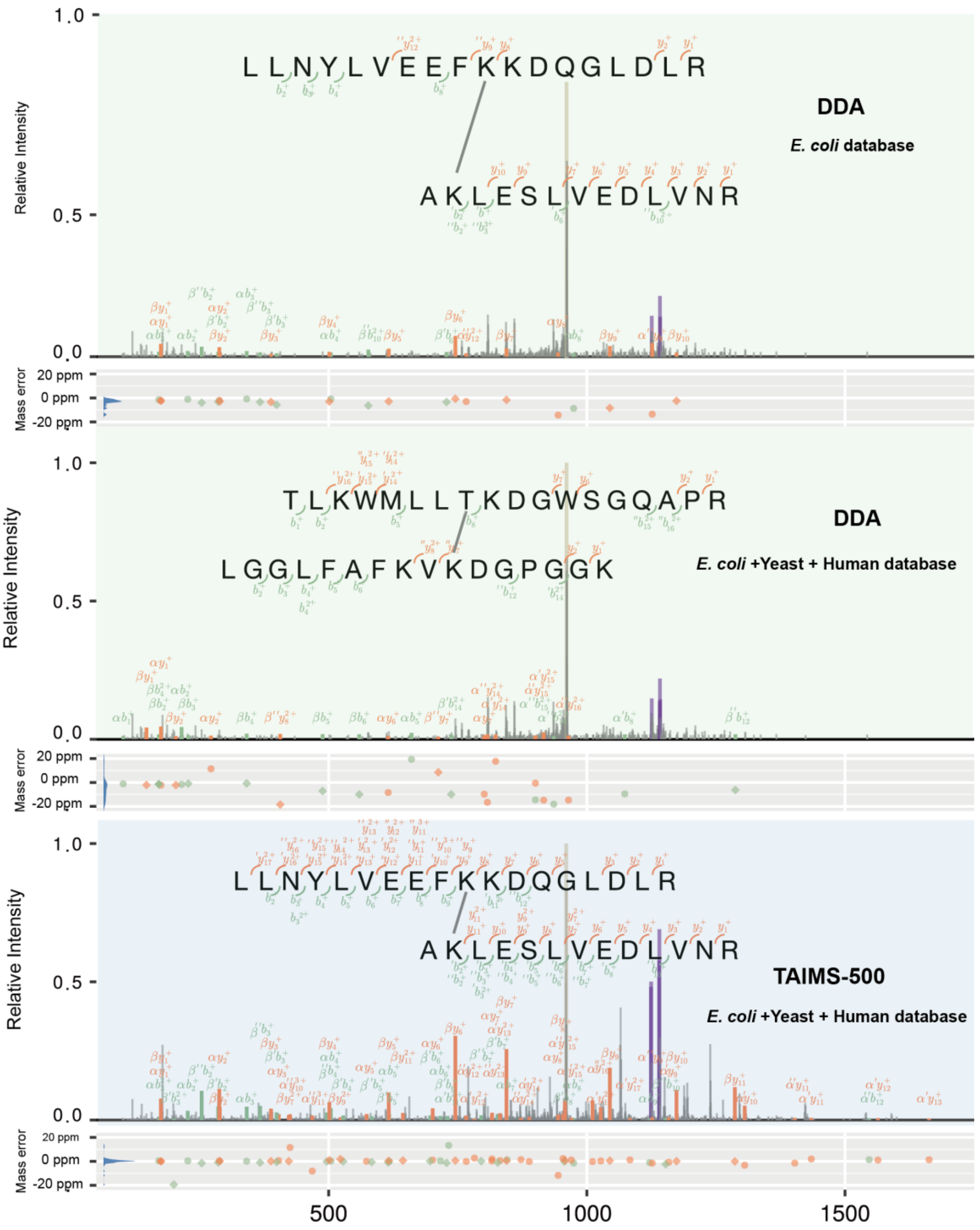


*Supplementary Figure 11. An example of TAIMS spectra resisting sensitivity loss in cross-link identification caused by search space expansion.*

The DDA MS2 of the indicated cross-link was mis-assigned to another cross-link in the expanded database search. In contrast, identification of the cognate TAIMS MS2 was not affected.

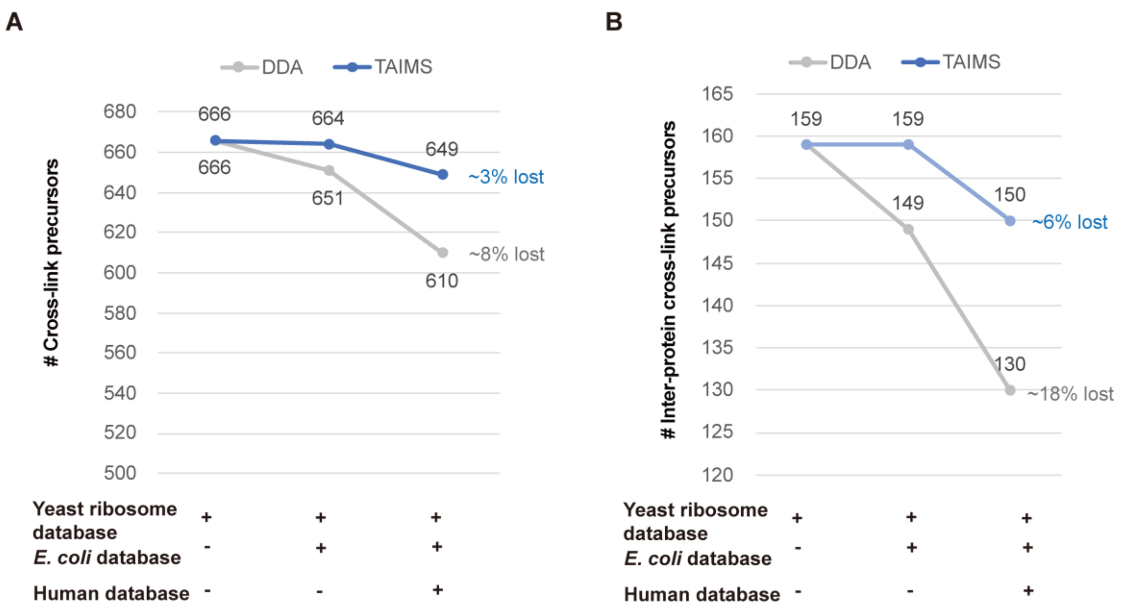


*Supplementary Figure 12. Statistics showing resistance of TAIMS spectra to sensitivity loss from search space expansion in a DSSO cross-linked yeast ribosome sample.*

The DDA and TAIMS data from the DSSO-cross-linked yeast ribosome sample yielded a common set of 666 total cross-link precursors (including 159 inter-protein cross-link precursors) when searched against the yeast ribosome database. The corresponding DDA spectra, but not the TAIMS spectra, suffered considerable sensitivity loss overall **(A)** and particularly for inter-protein cross-links **(B)** when searched against expanded databases that included *E. coli* and human databases in addition to the yeast ribosome database.

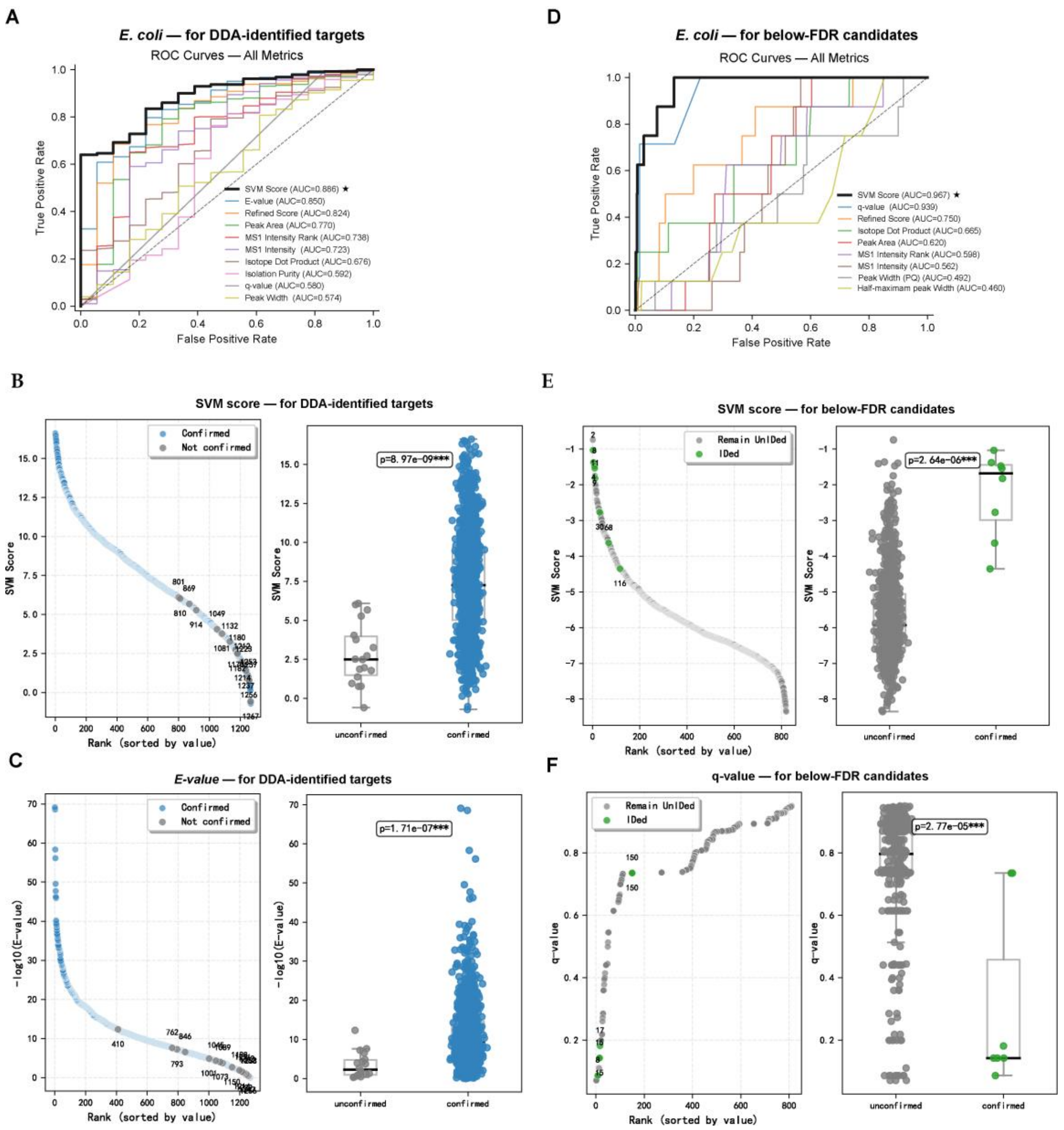


*Supplementary Figure 13. Prioritization of candidate targets for TAIMS analysis.*

**(A-C)** DDA-identified targets. (A) ROC curves comparing multiple quality metrics for their ability to distinguish TAIMS-confirmed (N = 1250) from TAIMS-unconfirmed (N = 18) precursors. True positive rate is calculated by counting the total number of DDA IDs above a given threshold that are later confirmed by TAIMS and dividing it by 1250. False positive rate is calculated by counting the total number of DDA IDs above a given threshold but are later unconfirmed by TAIMS and dividing it by 18. The score distributions of two best performing metrics, SVM score (AUC = 0.886) and *E*-value (AUC = 0.850), are shown in (B) and (C), respectively. **(D-F)** DDA-unidentified targets, i.e., CSMs below 5% FDR. (D) ROC curves comparing multiple quality metrics for their ability to distinguish TAIMS-salvaged (N = 8) from TAIMS-unsalvaged (N = 812) precursors. SVM score (AUC = 0.967) and q-value (AUC = 0.939) are the strongest discriminators. The distribution of SVM scores and that of q-values are displayed in (E) and (F), respectively.

Metric definitions: *E-value* (pLink2) is the expectation value for a given CSM; lower values indicate higher confidence. SVM Score (pLink2) is a support vector machine-based score distinguishing correct from incorrect CSMs; higher values indicate greater confidence. q-value (pLink2) is the minimum FDR at which a given precursor-level identification is accepted. Dot Product (Skyline) measures similarity between observed and theoretical isotopic distributions. Peak Area (Skyline) is the integrated area under the chromatographic peak. Half-width (Half-Maximum Peak Width, Skyline) is the chromatographic peak width at 50% maximum intensity. MS1 Intensity (pQuant) is the precursor ion signal intensity. MS1 Intensity Rank (computed from raw file) is the relative intensity rank among all MS1 precursors in the same scan. Isolation Purity (computed from MS1) is the fraction of target ion signal within the MS2 isolation window.

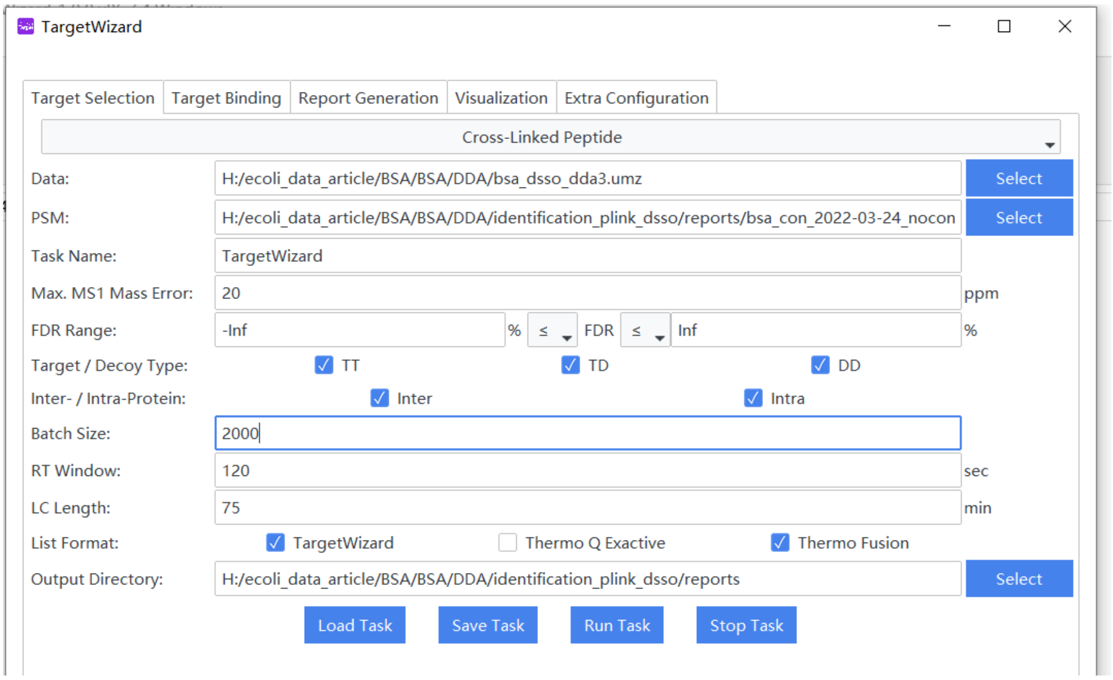


*Supplementary Figure 14. Multi-batch scheduling in TargetWizard.*

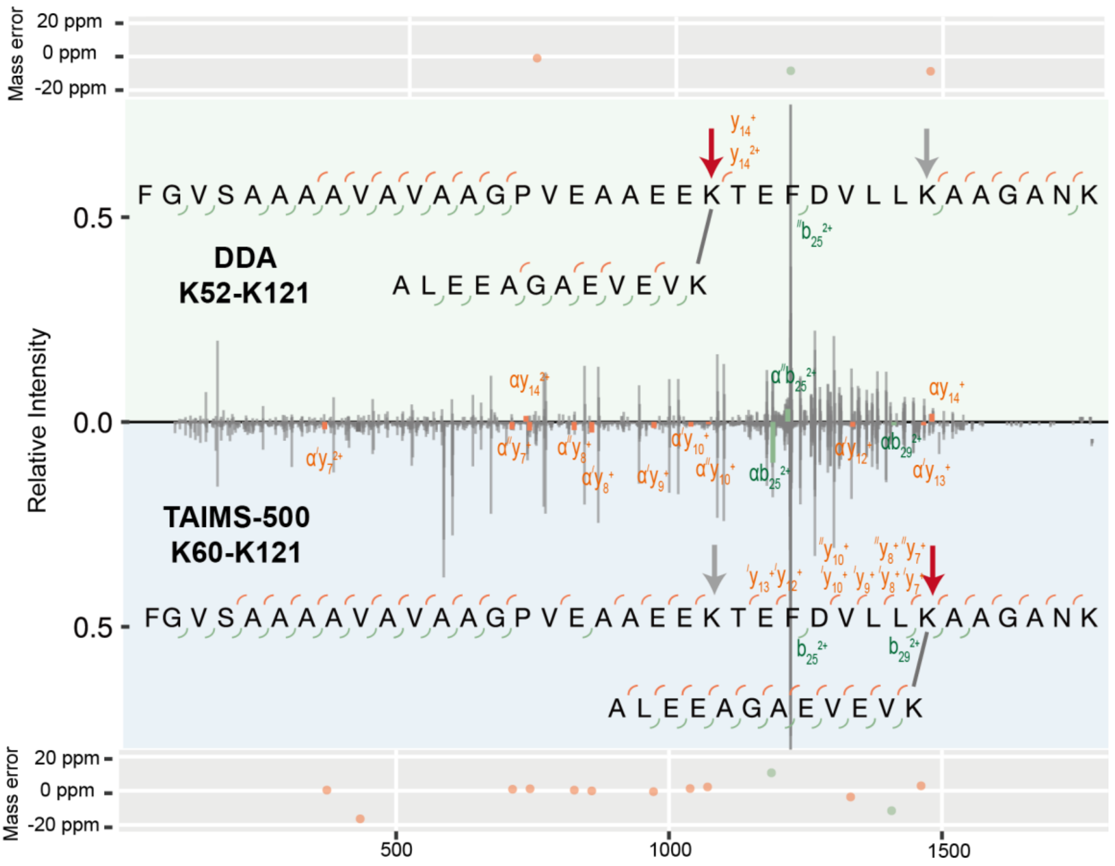


*Supplementary Figure 15. TAIMS improves the localization accuracy of cross-linked sites.*

DDA and TAIMS MS2 identified a cross-link between the same two peptides, both from protein L7/L12, but with different cross-linked sites in the α-peptide. For clarity, labeling of fragment ions is limited to the ones arising from a cleavage event between the two cross-link sites in question. Mass deviations of the fragment ions for locating the cross-linked residue in the alpha peptide are indicated at the top (for the DDA spectrum) or the bottom (for the TAIMS spectrum). For this cross-link, the cross-linked residue on the β-peptide is located at the C-terminus; this lysine is the last amino acid of the protein and thus does not violate the digestion specificity of trypsin.

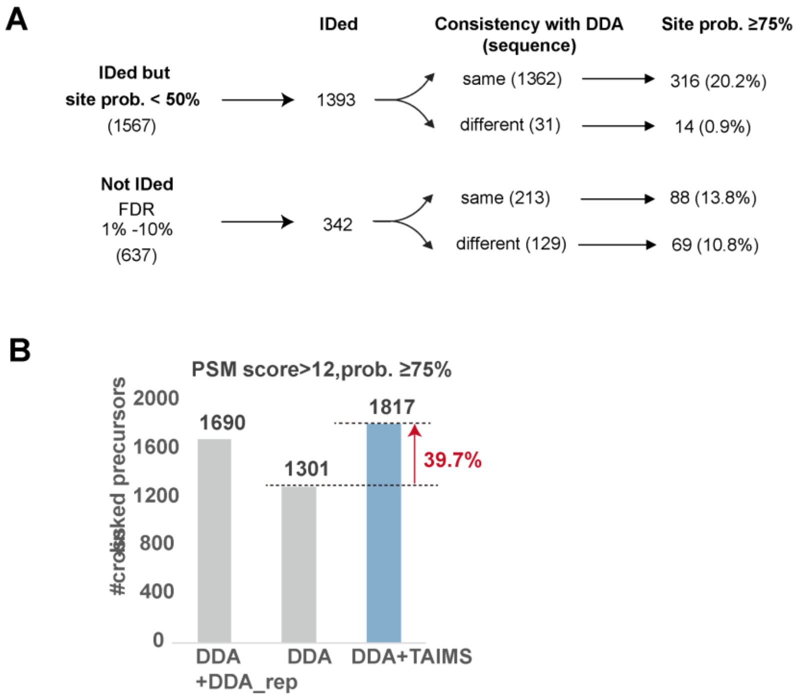


*Supplementary Figure 16. Phosphosite localization using a probability threshold of 75%.*

**(A)** TAIMS analysis result at 75% probability threshold. **(B)** Comparison of high-confidence phosphopeptide identifications (pFind3 PSM score >12 and localization probability ≥75%) between DDA and DDA-TAIMS.

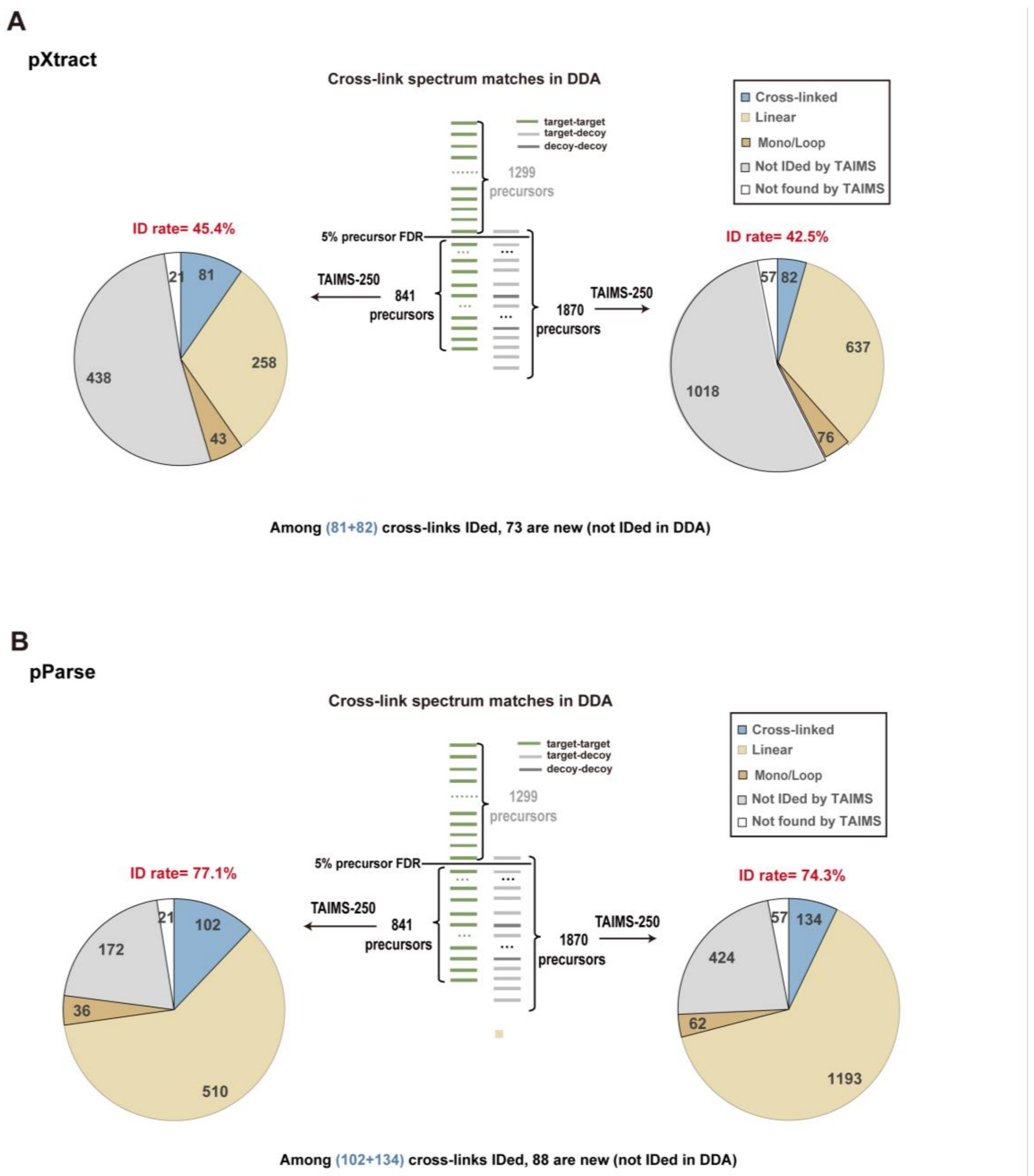


*Supplementary Figure 17. TAIMS analysis outcome of subthreshold DDA entries using preprocessing tools alternative to TargetWizard.*

The raw file of TAIMS data were preprocessed using **(A)** pXtract and **(B)** pParse. This analysis parallels what is shown in Figure 5A, where TargetWizard was used.

*Supplementary Table 1.* Recommendation on how to shorten an inclusion list for TAIMS

| Dataset | DDA Target Status | Total | TAIMS-ID | TAIMS-unID | Key Metrics (AUC) | Recommendation |
|---|---|---|---|---|---|---|
| *E. coli* | Identified | 1268 | 1250 | 18 | SVM (0.886) / E-value (0.850) | Filter by E-value $<1\times10^{-8}$ or $<1\times10^{-13}$ |
| | Unidentified | 820 | 8 | 812 | SVM (0.915) / E-value (0.875) | Filter by idotP ⩾ 0.85, rank by SVM, include top 20% |
| Yeast ribosome | Identified | 846 | 815 | 31 | SVM (0.967) / FDR (0.939) | Filter by E-value $<1\times10^{-9}$ or $<1\times10^{-13}$ |
| | Unidentified | 191 | 16 | 175 | SVM (0.785) / FDR (0.765) / idotP (0.712) | Filter idotP ⩾ 0.85, rank by SVM, include top 50% |

*Supplementary Table 2. Data analyses and datasets used in this study*

| Dataset name | Sample | Cross-linker | Mass spectrometry | #Raw Files |
|---|---|---|---|---|
| Optimization of MS2 AGC, MIT, and Resolution | synthetic peptides | DSSO | Fusion Lumos | 3 |
| Optimization of MIT and Dynamic Exclusion | BSA | DSSO | QE-HF | 10 |
| Optimization of MS2 MIT and Resolution | BSA | DSSO | QE-HF | 5 |
| Optimization of Global Intensity Threshold | *E. coli* lysates | DSSO | Fusion Lumos | 6 |
| Optimization of Individualized Intensity Threshold | yeast ribosome | DSSO | Fusion Lumos | 9 |
| Comparison of AIMS and PRM | yeast ribosome | DSSO | Fusion Lumos | 6 |
| Optimization of Dynamic Exclusion under Different Peak Widths | *E. coli* lysates | DSSO | Fusion Lumos | 21 |
| Evaluation of TAIMS Performance on Cross-linked Samples | *E. coli* lysates; yeast ribosome | DSSO | Fusion Lumos | 6 |
| Evaluation of TAIMS Performance on DSS Cross-linked Samples | BSA | DSS | Fusion Lumos | 2 |
| Evaluation of TAIMS Performance on pLink3 | yeast ribosome | DSSO | Fusion Lumos | 6 |
| Evaluation of TAIMS Performance on Phosphorylation | Mouse brain phosphopeptides | - | QE-HF | 3 |
